# The golden Syrian hamster (*Mesocricetus auratus*) as a model to decipher relevant pathogenic aspects of sheep-associated malignant catarrhal fever

**DOI:** 10.1101/2024.09.06.611580

**Authors:** Rosalie Fabian, Eleanor G Bentley, Adam Kirby, Parul Sharma, James P. Stewart, Anja Kipar

## Abstract

Malignant catarrhal fever (MCF) is an often fatal sporadic gammaherpesvirus-induced disease of ruminants with global relevance. Ovine gammaherpesvirus-2 (OvHV-2), with sheep as reservoir host, is a major cause of MCF in susceptible species. Despite extensive research on the molecular aspects of the disease, its pathogenesis is not yet fully understood. The present study re-established the Syrian golden hamster (*Mesocricetus auratus*) as amenable animal model of MCF and applied complementary *in situ* approaches to confirm recent findings in natural disease that could shed new light on pathogenetic aspects of MCF. These showed that systemic OvHV-2 infection is associated with T cell and macrophage dominated mononuclear infiltrates and vasculitis in various organs. Both T cells and monocytes/macrophages harbor the virus, and infected leukocytes are abundant in the infiltrates. The results also indicate that OvHV-2 has a broader target cell spectrum, including vascular endothelial cells and selected squamous epithelia. The former supports the interpretation that the inflammatory processes develop due to circulating activated, infected T cells and monocytes that home to tissues and emigrate from vessels prone to leukocyte emigration, possibly with direct interaction between virus infected leukocytes and endothelial cells. The latter supports the hypothesis of a graft versus host disease scenario, without viral cytopathic effect on epithelial cells but infiltration of the mucosa by infected T cells and macrophages. The disease processes are accompanied by evidence of expansion of the T cell compartments and the monocyte/macrophage pool in lymphatic tissues and bone marrow.

---

Malignant catarrhal fever (MCF) is a generally fatal disease caused by a group of ruminant gammaherpesviruses (subfamily *Gammaherpesvirinae*, genus Macavirus).^20^ Sheep associated MCF (SA-MCF), caused by ovine herpesvirus 2 (OvHV-2; formally *Macavirus ovinegamma2*),^17,20,60,72^ occurs worldwide in a wide range of susceptible hosts, including cattle, bison and deer, and occasionally pigs and goats.^2,11,39,42^ Alongside wildebeest-associated (WA-)MCF, SA-MCF is the most relevant form of the disease the clinical signs and pathological changes of which are similar regardless of the specific MCF virus involved.^61,76^

OvHV-2 is endemic in sheep which represent the natural reservoir host (so-called “host species”).^36^ Sheep are known as unaffected carriers of OvHV-2 and usually harbour the virus in circulating lymphocytes and/or the epithelium of the respiratory tract.^27,74^ However, they are not entirely resistant to SA-MCF and can develop MCF-like disease, both after natural infections and experimental exposure to very high viral inoculation doses.^18,36,37,55,56,74^

The pathological lesions in MCF are well known and similar in both cattle and bison, a species in which MCF has become of high economical relevance.^6,51^ However, while being extensively examined and reported, they are most comprehensively described in a widely read textbook.^22,51,52,66,76^ MCF is characterized by a combination of pathological processes; lymphoid hyperplasia, disseminated mononuclear vasculitis, and erosive/ulcerative mucosal lesions are the most relevant (Supplemental Table S1).^22,51,52,66,76^

It is widely accepted that T cells, and in particular cytotoxic T cells, play a major role in SA-MCF and dominate the inflammatory infiltrates observed in affected cattle,^44,61,63^ although some studies have shown an equal contribution of monocytes/macrophages.^49,63^ Similarly, circulating T cells were found to carry the virus in animals with MCF,^54,68^ but also in (experimentally) infected sheep where no restriction to one T cell subtype was detected (both CD2+CD4+ and CD2+CD8+) and even some monocytes (CD14+) were found to be infected.^45^ As part of a recent in-depth study on the *rete mirabile* vasculitis in bovine MCF, our group confirmed that virus infected T cells and monocytes are both participating in the process.^63^

So far, despite extensive research on the molecular aspects of the disease and its variable clinical presentation, the pathogenesis of MCF is not fully understood. Based on the recent knowledge, it is hypothesised that the pathological processes in MCF are based on an abnormal cytotoxic T cell activity initiating a graft-versus-host like reaction that attacks epithelia and arteries.^76^ Interestingly, the concept of a graft-versus-host like pathogenesis was already established before the causative virus was identified, more than 40 years ago.^40^

In natural MCF virus infections, the upper respiratory tract and/or the tonsils are the most likely portal of entry, and this route of infection has also been used in experimental settings with large animals and rabbits. Most common practice is to harvest virus from infected animals for subsequent experimental infection. Animals can become infected with OvHV-2 when nebulized with pooled infected sheep nasal secretion containing defined high DNA copy numbers (10^7^ DNA copies).^16,34^ Inoculation with infected lymphoid cells is an alternative option and can use the intravenous and intraperitoneal route.^11,16,33,34,57,65^ *In vitro* propagation of OvHV-2 is only possible in the established lymphocyte cell line BJ1035, an immortalized anergic T cell line originating from a clinical cattle MCF case.^4,7,13,20,46,58,64,65,71^ Such studies provided much of what is currently known about OvHV-2 infection and disease development in deer, cattle and bison.^13^

Due to practicality and affordability, small animal models allow for more in-depth research into the pathogenesis of the disease and modulation and monitoring of host immune response which likely play a relevant role for susceptibility and development of MCF.^8,35^ Since the first reports in the late 1980s,^9,10^ the rabbit has been established and progressively promoted as a model for SA- and WA-MCF, in recent years also for vaccine development.^12–14,65^

Pathological descriptions of the disease in rabbits vary in detail but consistent changes include: lymphocyte/lymphoid cell/T cell infiltrations in liver (portal areas), lungs (perivascular, -bronchial, -bronchiolar, interstitial) and kidneys (perivascular, interstitial), and arteritis-phlebitis in liver and lungs; similar infiltrates are reported as occasional in various organs/tissues;^4,13,16,34,65^ necrotic or granulomatous hepatitis and lymphadenitis have also been observed.^13,51^ Lymphatic tissues (lymph nodes, spleen, appendix) are described as hyperplastic, affecting the T cell zones.^4,10,16,34,65^ Ulcerative lesions were reported on tongue and cecum,^13^ as well as subepithelial infiltrates in the esophagus.^65^ The brains and eyes have either not been examined histologically or were found to be unaffected.^13^ The reported findings are summarized in Supplemental Table S1.

Whilst the rabbit model reflects natural MCF in many aspects, it is relatively resistant to OvHV-2 infection and requires a lethal dose 2-fold higher than, for example, a bison.^15,67^

The hamster has been used as an animal model to study human diseases for over 60 years.^47^ It has served for experimental infections with more than 70 different viruses, among these paramyxoviridae and filoviridae, phlebo- and flaviviruses as well as arenaviridae, with a rising trend.^47^ In particular during the last 4 years, hundreds of studies used the hamster as model for COVID-19. The hamster has also been proposed as suitable to model MCF.^8,30,57^ An initial experiment published in 1981 reports inoculation of newborn hamsters with 5×10^3^ TCID50 ‘malignant catarrhal disease virus’ isolated from African wildebeest calves with MCF.^30^ This was followed a few years later by a more thorough examination where, after intraperitoneal inoculation with lymph node cells isolated from a red deer and a cow with MCF that were passaged several times in hamsters;^8,28^ animals became clinically ill and developed histological changes that appeared in a broader sense similar to those described in cattle with MCF.^8^ Since then, this animal model has not been pursued further.

The present study aimed to re-establish the hamster model of MCF and extend the pathological observations using modern molecular techniques. To do this, Syrian golden hamsters (*Mesocricetus auratus*) were intraperitoneally inoculated with OvHV-2 carrying lymphoid cells from a hamster, clinically monitored and euthanized at clinical endpoints. After confirmation of systemic infection by PCR, histological and immunohistochemical examinations and RNA *in situ* hybridization served to characterize the pathological processes and identify viral target cells.

## Material and Methods

### Cell culture and virus

In an initial experiment, ovine herpesvirus-2 (OvHV-2) infected hamster lymphoid cells were kindly provided by Dr George Russell (Moredun Institute, Edinburgh, UK). The cells were from a previous experiment undertaken in 2005 where hamsters were experimentally infected with sheep-associated OvHV-2 and euthanised when they presented with either loss of appetite, depression, diarrhea and/or ocular/nasal discharge. From the hamsters of this pilot experiment, spleen and lymph nodes were collected and passed through a 70 µm cell strainer (Corning, Corning, USA), and the cells collected were stored in liquid nitrogen until their use in the following larger experiment.

Prior to infection, cells were raised from liquid nitrogen then live dead sorted using the EasySep™ Dead Cell Removal (Annexin V) Kit (Stemcell Technologies, Vancouver, Canada) according to the manufacturer’s instruction. Live cells were adjusted to 1×10^6^ cells/ml and stored on ice ready to use.

### Hamsters

Animal work was approved by the local University of Liverpool Animal Welfare and Ethical Review Body and performed under UK Home Office Project Licence PP4715265. Male 8-10 week old golden Syrian hamsters were purchased from Janvier Labs (France). Hamsters were maintained under SPF barrier conditions in individually ventilated cages and fed pelleted food *ad libitum* with free access to water. Prior to experimentation hamsters were acclimatised to their cage for 1 week.

### Experimental infections

In both the initial study (n=4) and the subsequent main study (n=6), animals were inoculated intraperitoneally with pooled spleen and lymph node cells from an OvHV-2 infected hamster (see above). In the first experiment, each animal received 2.2×10^5 cells which was estimated to be 1.5×10^8 copies/dose. In the second experiment, the animals were inoculated with 1.3×10^5 cells per hamster which was estimated to be 1.2×10^6 copies/dose.

All animals were closely monitored after inoculation for adverse effects including weight loss, hunching, isolation and ruffled fur. Body temperature was monitored daily with a subcutaneous temperature transponder (Plexx, Netherlands) and health condition were monitored daily.

Infection was defined as established when an animal sustained an increase of 1 °C in its average body temperature (defined as the average of the first 5 days recorded) for over 48 h.^46^ Other clinical signs included but were not limited to facial grimace, scruffy coat, diarrhea and swollen feet. Once infection was clinically confirmed animals were euthanised by a schedule 1 method and immediately dissected after death.

For the animals in the initial study, blood samples were collected under terminal anesthesia into EDTA containing vacutainers (Becton Dickson, Wokingham, UK) and centrifuged at 800 *x g* for 10 min. The buffy coat was collected for DNA extraction and quantitative PCR. A selection of organs suspected to exhibit pathological changes based on the clinical signs was collected and fixed in 10% buffered formalin. From the animals in the main experiment, all major organs as well as the entire body were fixed in 10% buffered formalin for histological examination.

### DNA extraction and quantification

Buffy coat cells were homogenised and DNA extracted using the DNeasy Blood & Tissue Kit (Qiagen, Manchster, UK) according to the manufacturer’s instructions. The extracted DNA was quantified using a Nanodrop one spectrophotometer (Thermo Fisher, Waltham, US).

### Quantitative polymerase chain reaction (qPCR) for OvHV-2

For qPCR, TaqMan^TM^ Universal PCR Master Mix (Applied Biosystems^TM^) was used in 20 µl reaction volumes. Each mix consisted of 10 µl TaqMan® Universal PCR Master Mix (2X), 2 µl each of forward and reverse primers (100 nM) and 2 µl probe (250 nM) for either OvHV-2 or 18s-ribosomal DNA and 100 ng template DNA suspended in 4 µl nuclease-free water. OvHV-2 and 18s primers were as described in previous studies.^59,69^ Detection of the 18s-ribosomal DNA internal genome served to normalize for DNA variability and contaminants to give viral copies/µg. The RT-PCR was run with the following cycling conditions: 50 °C for 2 min, 95 °C for 10 min, then 39 cycles of 95 °C for 15 sec and 60 °C for 1 min. The data were collected during step 4 of the PCR program.

### Histology, immunohistology and RNA in situ hybridization

From animals in the initial study, a selection of organs (skin, tongue, stomach, small intestine, liver, spleen, lungs, kidneys, and the brain from 3 animals, lung from the fourth) was processed. From all animals in the main experiment, samples from tongue, esophagus, stomach, small intestine, liver, spleen, lymph nodes (cervical, mediastinal, mesenteric), heart, trachea, lungs, kidneys, brain, lumbar spinal cord (cross section and longitudinal section of 3-5 mm length), right sciatic nerve and the *M. biceps femoris* were processed. The tissue samples were trimmed and routinely embedded in paraffin wax. Consecutive sections (3-4 μm thick) were prepared and routinely stained with hematoxylin and eosin (HE) and, in selected specimens, subjected to immunohistology (IH) and RNA *in situ* hybridization (RNA-ISH). Lymphatic tissues (mediastinal lymph nodes, spleen) from age-matched mock infected control hamsters of an unrelated COVID-19 study served for the comparative assessment with the tissues from the infected hamsters in the present study.

IH served to characterize the infiltrating leukocytes, i.e. T cells (CD3+), B cells (CD79a+), macrophages (Iba-1+), and highlight proliferating cells (PCNA+) and apoptotic cells (cleaved caspase 3+) using cross-reactive antibodies and previously established protocols.^23,26,43,53,70^ Stains were carried out in an autostainer (“Link 48” (Agilent Dako), using the horseradish peroxidase method. Briefly, sections were deparaffinised through graded alcohol and subjected to antigen retrieval in citrate buffer (pH 6.0) or Tris/EDTA buffer (pH 9) for 20 min at 98 °C. Slides were subsequently incubated with the primary antibodies diluted in antibody diluent (Dako). This was followed by blocking of endogenous peroxidase (peroxidase block, Dako) for 10 min at room temperature and incubation with the appropriate secondary antibodies/detection systems following the manufacturers’ protocols. All antibodies and detection methods are listed in Supplemental Table S2. Sections were washed with phosphate buffered saline (pH 8) between each incubation step. They were counterstained with hematoxylin for 20 s and mounted. The lymph node from an uninfected control hamster served as a positive control for all markers, sections incubated without the primary antibodies served as negative controls.

RNA-ISH was performed using the RNAscope® ISH method (Advanced Cell Diagnostics (ACD, Newark, California), Ov2.5 mRNA (coding for OvHV-2 viral IL-10; Genbank NC_007646.1) oligoprobes and the automated RNAscope® 2.5 Detection Reagent Kit (Brown) according to the manufacturer’s protocol, and as previously published, applying slight modifications.^42,63^ Briefly, sections were heated to 60 °C for 1 h, deparaffinized and permeabilized by incubation in pretreatment solution 1 (RNAscope^®^ Hydrogen Peroxide) for 10 min at RT, followed by boiling in RNAscope^®^ 1X Target Retrieval Reagents solution at 100 °C for 25 min and washing in distilled water and ethanol. This was followed by digestion with RNAscope^®^ Protease Plus for 30 min and hybridization with the oligoprobes for 2 h, and serial amplification with different amplifying solutions (AMP1, AMP2, AMP3, AMP4: alternating 15 min and 30 min), all at 40 °C in a humidity control tray for 2 h (HybEZ^TM^ Oven, ACD Advanced Cell Diagnostics). Between each incubation step, slides were washed for 2 x 2 min with washing buffer. They were subsequently incubated with AMP 5 (45 min), AMP 6 (15 min) and DAB (10 min) at RT in the humidity control tray. Sections were counterstained for 15 s with Gill’s hematoxylin, then dehydrated with graded alcohol and xylene and coverslipped. A formalin-fixed, paraffin-embedded cell pellet of BJ1035 cells (permanently OvHV-2 infected bovine large granular lymphocyte cell line) served as a positive control.^63^ Consecutive sections incubated accordingly but without including the hybridization step and sections from a formalin-fixed, paraffin-embedded liver sample of a hamster not infected with OvHV-2 served as negative controls.

In order to determine the leukocytes that carry the virus, a combined RNA-ISH/IH protocol was applied to sections of the liver of animal No. 10. As a first step, IH for CD3 and Iba1 was undertaken after the pretreatment steps for the ISH (baking, deparaffinizing, incubating in RNAscope^®^ Hydrogen Peroxide, cooking in RNAscope^®^ 1X Target Retrieval Reagents solution and incubating in RNAscope^®^ Protease Plus) instead of the normal antigen retrieval process for IH. Since the IH staining was maintained after this pretreatment, further sections were subjected to the entire RNA-ISH protocol, without letting the sections dry at any time point during the procedure.

Subsequently, the sections were incubated in tap water instead of counterstaining with hematoxylin and dehydration, followed by the respective IH protocol, with overnight incubation of the antibodies at the dilutions indicated in Supplemental Table S2. The reaction was visualized with the same kit as indicated in Supplemental Table S2 but with AEC single solution (Zytomed Systems GmbH, Berlin, Germany) instead of DAB, followed by counterstaining for IH.

## Results

### Intraperitoneal infection of golden Syrian hamsters with OvHV-2 leads to viraemia and clinical disease

In an initial study, four hamsters were inoculated intraperitoneally with spleen and lymph node cells derived from infected hamsters. Here, the first hamster (animal No 1) had shown sustained elevated temperature and further clinical signs (swollen left hind limb, limping) by 29 days post inoculation (dpi), the second appeared healthy but was euthanized at 33 dpi due to elevated body temperature, and the remaining two hamsters (animals No 3 and 4) were euthanized at 41 dpi for the same reason (Supplemental Table S3). One had also shown facial grimace and scruffy coat, the other diarrhea. In all four animals, the buffy coat was positive for OvHV-2 DNA by qPCR, with 94 to 1.5×10^5^ copies/µg, confirming that the animals were viremic at the time of death.

In the second experiment (six hamsters), the clinical course was more rapid. Animals were euthanized 15-17 dpi following a rise in body temperature and presentation of general malaise and a combination of other clinical signs including piloerection and facial grimace (Supplemental Table S3).

The post mortem examination did not reveal any gross changes in any animal, apart from evidence of a mild subcutaneous edema in the hamster with the swollen hindleg (animal no 1) and the presence of coagulated blood in the urinary bladder of animal no 6.

### Systemic OvHV-2 infection is associated with T cell and macrophage dominated mononuclear infiltrates in various organs and tissues as well as vasculitis, all with abundant OvHV-2 infected leukocytes

Animals in both experiments exhibited widely consistent changes in a range of organs, although these often varied in their extent. Information on affected organs is provided in Supplemental Table S3. Subsequently, the main findings are reported.

#### Liver (n=9)

All animals exhibited portal, predominantly mononuclear infiltrates, mostly of a moderate degree (consistent with moderate chronic hepatitis) as well as an increase in leukocytes within the sinusoids (Fig. 1). The infiltrates were comprised of T cells (CD3+) and, even more numerous, macrophages (Iba1+) of which many were positive for viral RNA (Ov2.5) (Fig. 1c-e). The same applied to the sinusoidal leukocytes (Fig. 1f-h). In some animals (nos 7, 8 and 10), the inflammatory infiltrates were associated with subendothelial leukocyte aggregation and focal infiltration of the wall in portal veins (Fig. 1b-e), consistent with vasculitis. One hamster (no 3) exhibited small endothelium-attached leukocyte thrombi in one portal vein.

**Figure 1.**
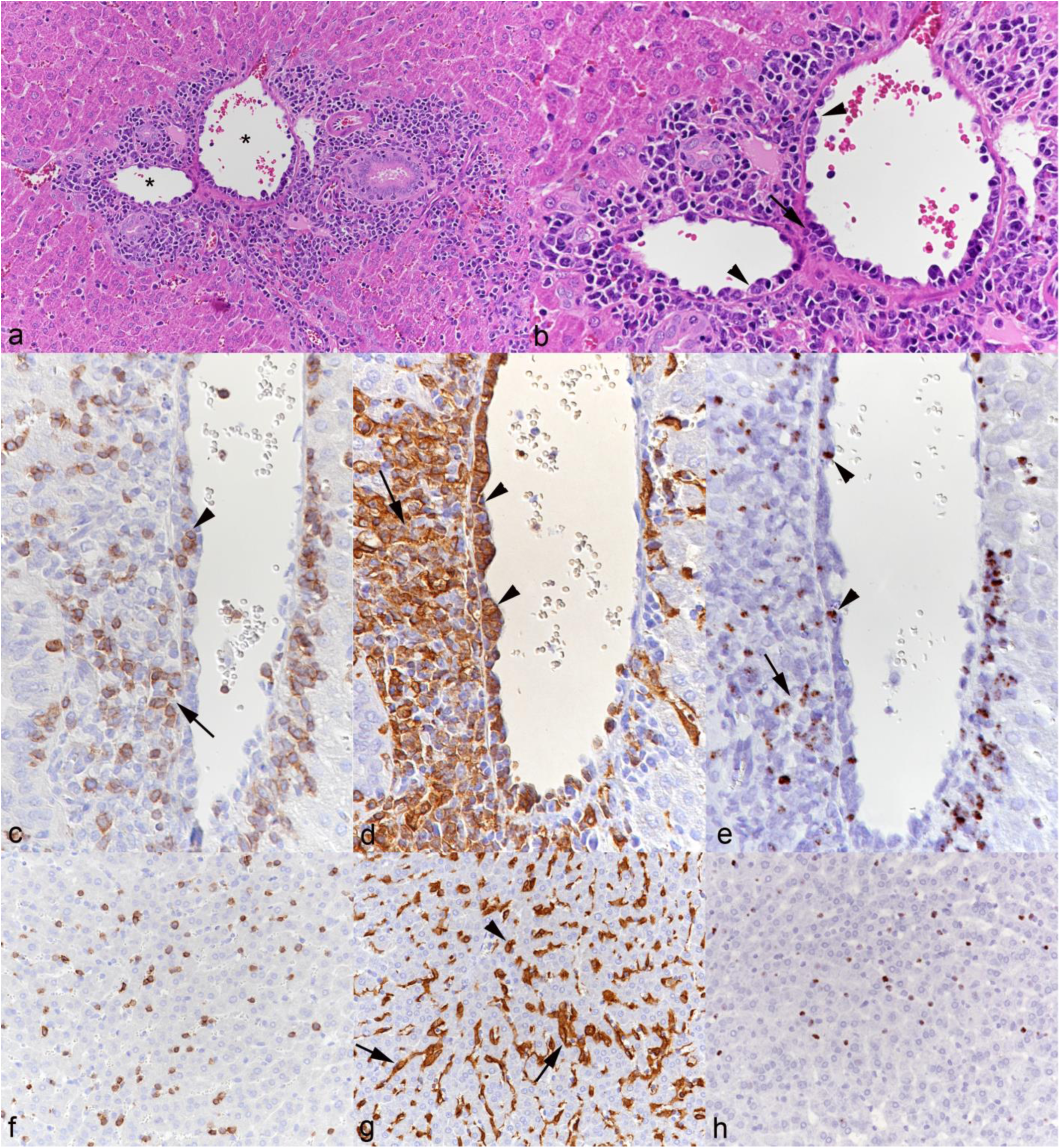
Liver, animal No. 10. **a.** Portal area with moderate mononuclear infiltration. Asterisk: portal veins with evidence of vasculitis. HE stain. **b.** Closer view of portal veins with subendothelial aggregations of leukocytes (arrowheads) and infiltration of the vessel wall (arrow) consistent with vasculitis. HE stain. **c-e.** Portal vein. **c.** Among the leukocytes in the subendothelial aggregates (arrowhead) and surrounding the vein are numerous T cells (CD3+). Immunohistochemistry (IH), hematoxylin counterstain. **d.** The subendothelial (arrowheads) and perivascular (arrow) leukocyte aggregates are dominated by monocytes/macrophages (Iba1+). IH, hematoxylin counterstain. **e.** Among the leukocytes in the subendothelial aggregates (arrowhead) and surrounding the vein (arrow) are several positive for viral RNA (Ov2.5). RNA-ISH, hematoxylin counterstain. **f-h.** Sinuses. **f.** Sinuses contain increased numbers of T cells (CD3+). IH, hematoxylin counterstain. **g.** Staining for Iba1 shows that besides Kupffer cells (arrows) the sinuses contain increased numbers of monocytes (arrowhead). IH, hematoxylin counterstain. **h.** Numerous leukocytes in the sinuses are positive for viral RNA (Ov2.5). RNA-ISH, hematoxylin counterstain.

#### Heart (n=6)

The myocardium and left atrioventricular valves were examined in animals in the main experiment (nos 5-10); these all exhibited a mild to moderate mononuclear valvular endocarditis, with clear subendothelial aggregation of leukocytes (T cells and monocytes in comparable proportions) many of which were positive for viral RNA (Fig. 2). In two hamsters (nos 7,10), the ventricular endocardium exhibited further identical infiltrates. In all animals, the myocardium showed mild multifocal interstitial, partially perivascular T cell and macrophage infiltrates.

**Figure 2.**
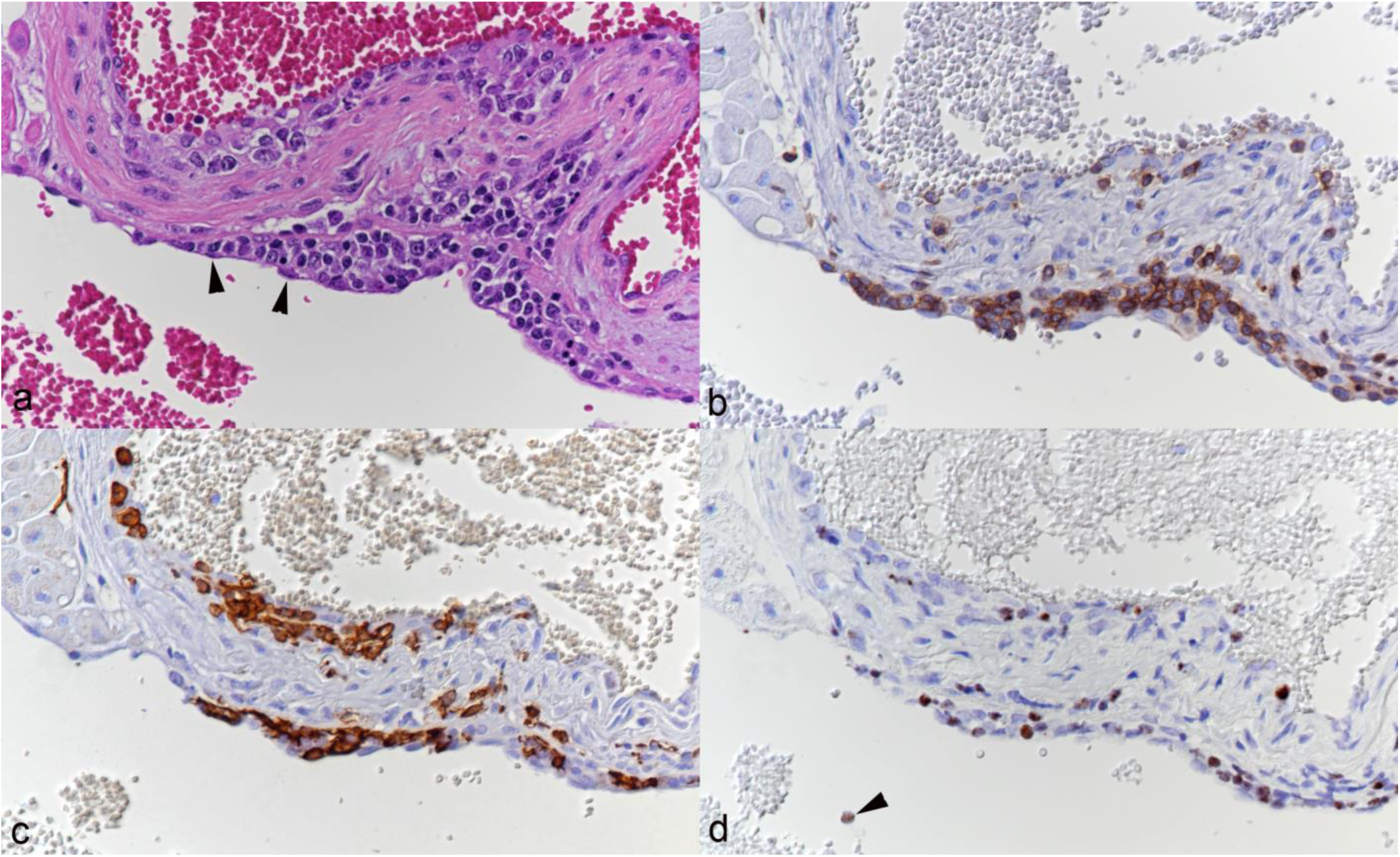
Heart, left atrioventricular valve, animal No. 8. **a.** Multifocal leukocyte infiltration beneath the endothelium (arrowheads). HE stain. **b.** Among the infiltrating leukocytes are abundant T cells (CD3+). Immunohistochemistry (IH), hematoxylin counterstain. **c.** Among the infiltrating leukocytes are abundant monocytes/macrophages (Iba1+). IH, hematoxylin counterstain. **d.** Several infiltrating leukocytes are positive for viral RNA (Ov2.5). Arrowhead: Ov2.5 mRNA positive leukocyte in the blood. RNA-ISH, hematoxylin counterstain.

#### Lungs (n=10)

The most consistent finding in the lungs of all animals was a substantial increase in T cells and, though less numerous, monocytes within capillaries (Fig. 3a, b); whether a proportion of the cells seen in the alveolar walls were located in the interstitium could not be clearly discerned. Many of the leukocytes were found to harbor viral RNA (Fig. 3c). B cells (CD79a+) were rare in capillaries. In a few animals there was evidence of leukocyte rolling and emigration (nos 1, 3, 4) or mild perivascular leukocyte infiltration (nos 1-5). One hamster (no 3) exhibited mild inflammation of an artery (Fig. 3d).

**Figure 3.**
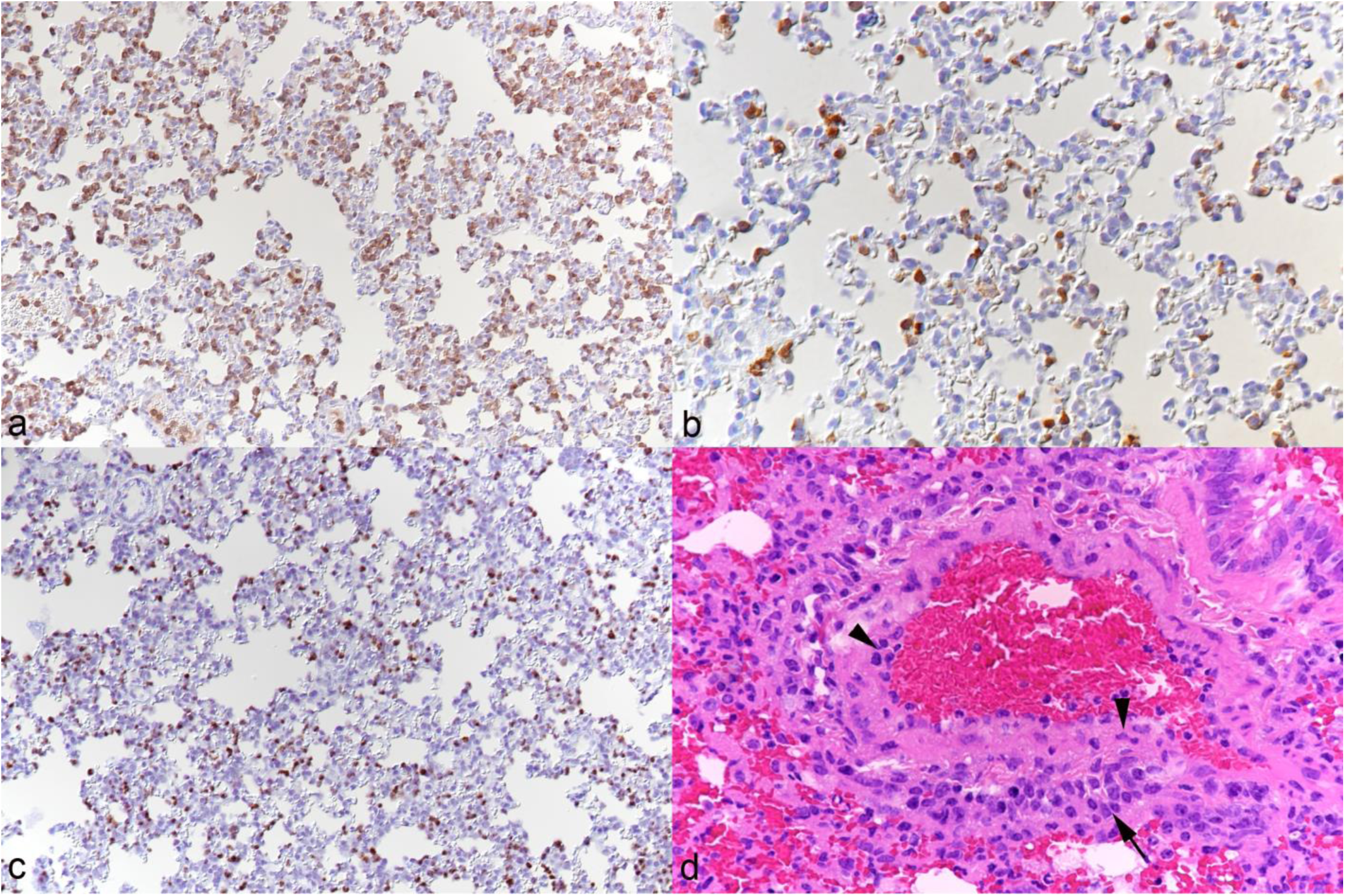
Lung. **a-c.** Animal No. 8. **a.** There are very abundant T cells (CD3+) within capillaries. Immunohistochemistry (IH), hematoxylin counterstain. **b.** Capillaries contain increased numbers of monocytes (Iba1+). IH, hematoxylin counterstain. **c.** Abundant leukocytes within capillaries are positive for viral RNA (Ov2.5). RNA-ISH, hematoxylin counterstain. **d.** Animal No 3. Artery with leukocytes in wall (arrowheads) and in the surrounding tissue (arrow), consistent with arteritis. HE stain.

#### Alimentary tract

The <u>tongue</u> (n=9) consistently exhibited mononuclear subepithelial infiltrates of varying intensity (Fig. 4). The infiltrate was comprised of T cells and macrophages which were also seen to invade the epithelium (Fig. 4b, c). In some cases, this was associated with apoptosis of individual epithelial cells (Fig. 4a). With more intense infiltration, inflammatory cells also stretched between the musculature. A proportion of infiltrating leukocytes were found to express viral RNA (Fig. 4d). In the <u>esophagus</u> (n=6), similar subepithelial infiltrates were observed. However, these were only focal and mild, with occasional individual leukocytes between basal epithelial cells.

**Figure 4.**
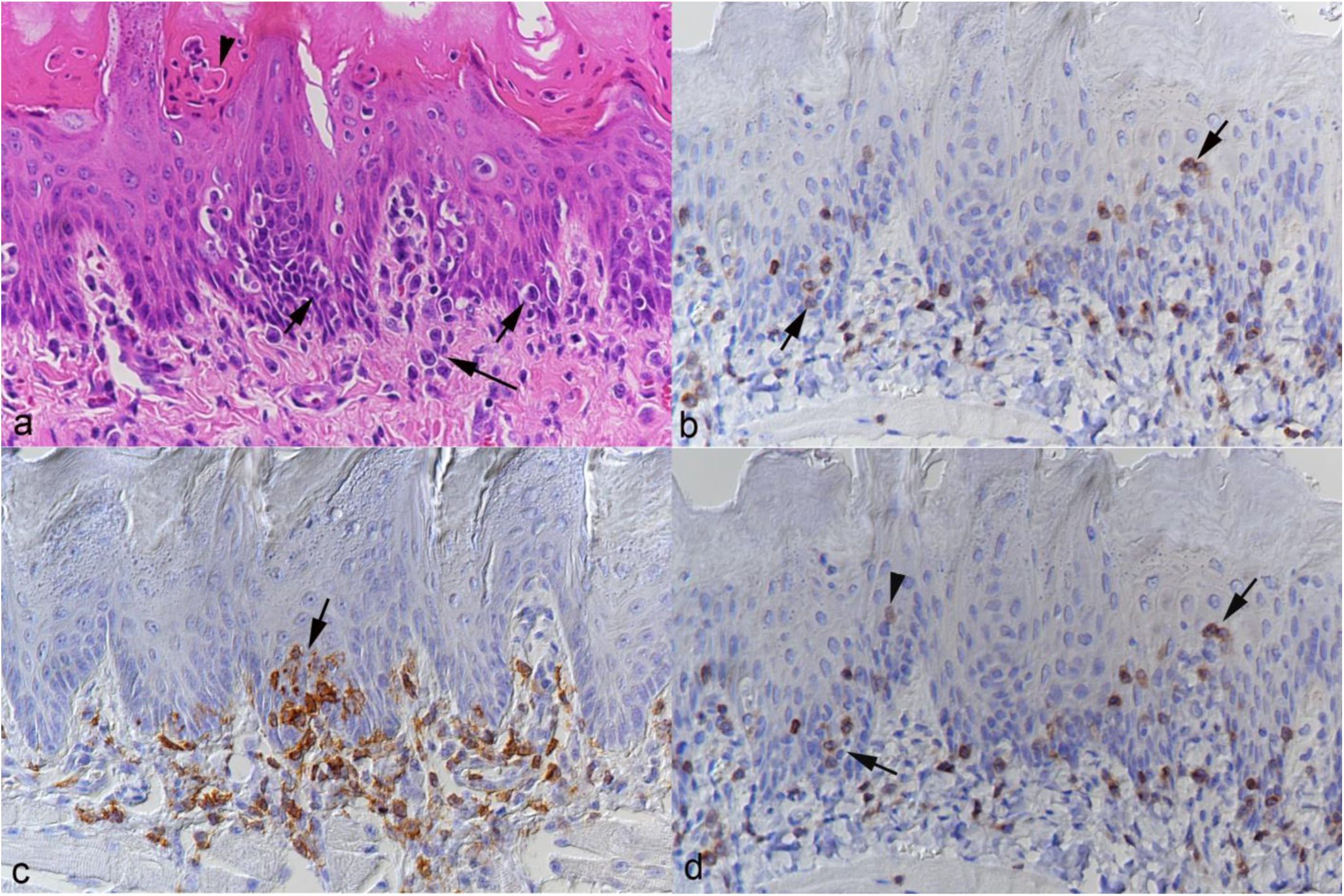
Tongue, animal No. 3. **a.** Leukocyte infiltration beneath (long arrow) and within (short arrows) the epithelial layer. There are also individual apoptotic epithelial cells (arrowhead). HE stain. **b.** Infiltrating T cells (CD3+) are found beneath the epithelium but are also present between epithelial cells (arrows). Immunohistochemistry (IH), hematoxylin counterstain. **c.** Focal macrophage (Iba1+) rich subepithelial infiltrate with aggregates of macrophages between epithelial cells (arrow). IH, hematoxylin counterstain. **d.** Abundant infiltrating leukocytes both subepithelially and infiltrating the epithelium (arrows) are positive for viral RNA (Ov2.5); there are also individual cells with a nuclear signal that have the morphology of epithelial cells (arrowhead). RNA-ISH, hematoxylin counterstain.

In most animals (nos 2-4, 7-10) the <u>stomach</u> (n=9) exhibited mild to moderate diffuse mononuclear infiltration of the mucosa; in three animals (nos 2-4), this stretched into the submucosa. Again, the infiltrates were comprised of T cells and macrophages, with a large proportion of cells positive for viral RNA. In the <u>non-glandular forestomach</u>, infiltrating cells were observed between the epithelial cells in three hamsters (nos 5, 6 and 10). Animal no 3 showed a severe erosive to ulcerative proventricultis with multifocal serocellular crust formation, pustule formation and abundant apoptotic keratinocytes (Fig. 5a, b; Supplemental Fig. S1). There was a marked mononuclear T cell and macrophage dominated submucosal infiltration that also invaded the basal epithelial layer and harbored abundant virus RNA positive leukocytes (Fig. 5c-f), with clear evidence of recruitment of infected leukocytes from the blood (Fig. 5f).

**Figure 5.**
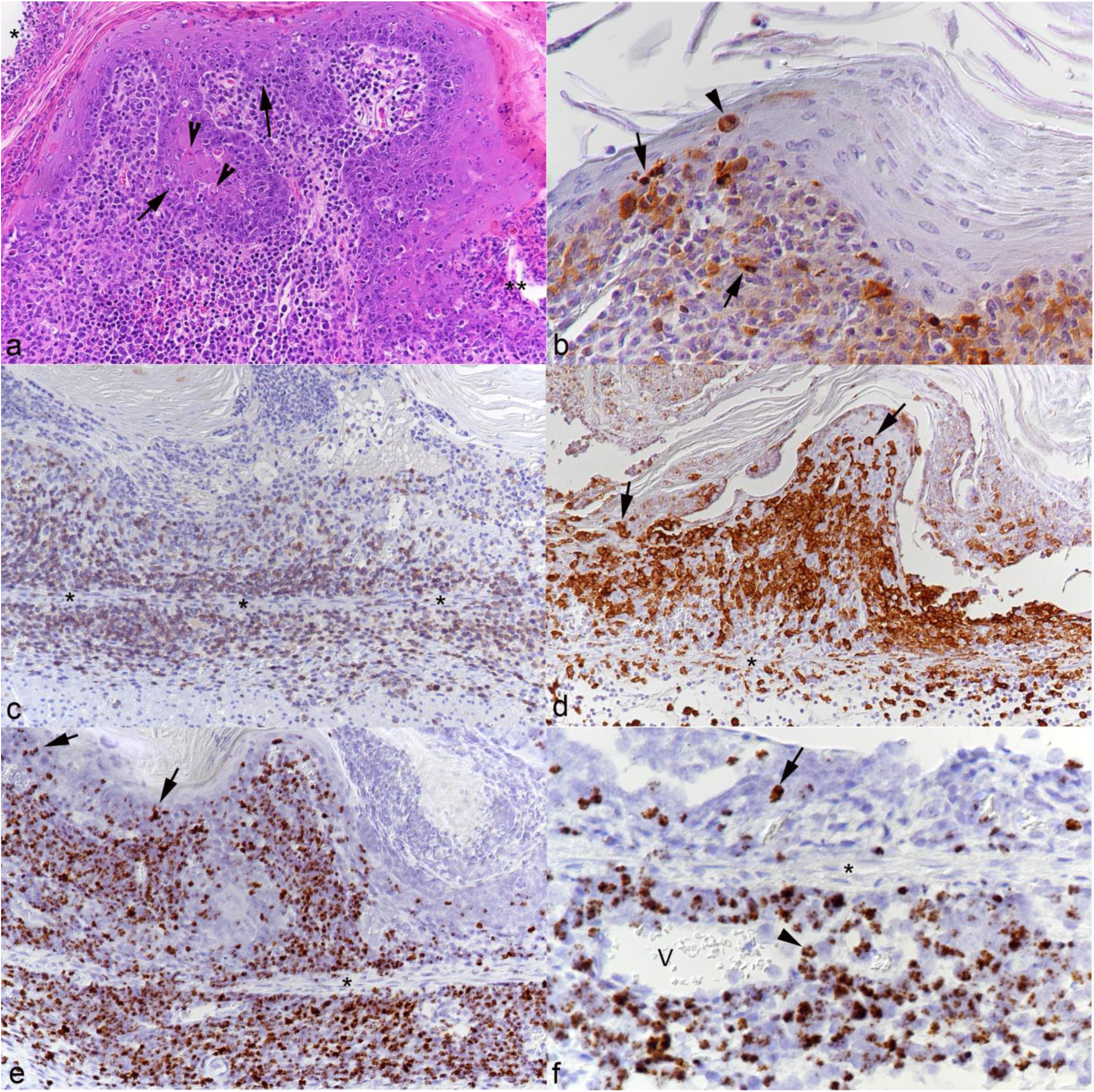
Non-glandular forestomach, animal No. 3. **a.** Serocellular crust formation (*) with focal ulceration (**) and marked subepithelial leukocyte infiltration. Leukocytes focally invade the basal epithelial cell layers (arrows). There are individual apoptotic epithelial cells (arrowheads). HE stain. **b.** Staining for cleaved caspase 3 confirms apoptosis of individual epithelial cells (arrowhead) and also shows that some leukocytes both within the epithelium and in the submucosal infiltrate undergo apoptosis (arrows). Immunohistochemistry (IH), hematoxylin counterstain. **c.** A large proportion of infiltrating leukocytes in the mucosa and submucosa are T cells (CD3+). The asterisks highlight the muscularis mucosae. IH, hematoxylin counterstain. **d.** Area with extensive serocellular crust formation. There are abundant macrophages (Iba1+) in particular in the mucosal infiltrate, also infiltrating the epithelium (arrows). The asterisk highlights the muscularis mucosae. IH, hematoxylin counterstain. **e, f.** RNA-ISH for Ov2.5, hematoxylin counterstain. The asterisks highlight the muscularis mucosae. **e.** Abundant leukocytes in the mucosal and submucosal infiltrates and infiltrating the epithelium (arrows) are positive for viral RNA. **f.** A closer view of a submucosal vein (V) highlights recruitment of viral RNA positive leukocytes (arrowhead). The epithelium contains a cell with a nuclear signal that has the morphology of an epithelial cell (arrow).

In all animals, the <u>small intestine</u> showed diffuse expansion of the villi due to a marked mononuclear infiltration dominated by T cells, followed by macrophages (Fig. 6a-c), with only rare B cells and plasma cells (Supplemental Fig. S2a). Abundant cells in the infiltrate were found to carry viral RNA (Fig. 6d); the latter was also seen in lymphocytes in the generally prominent peyer’s patches. The infiltration was often association with villus blunting and fusion (Supplemental Fig. S2b).

**Figure 6.**
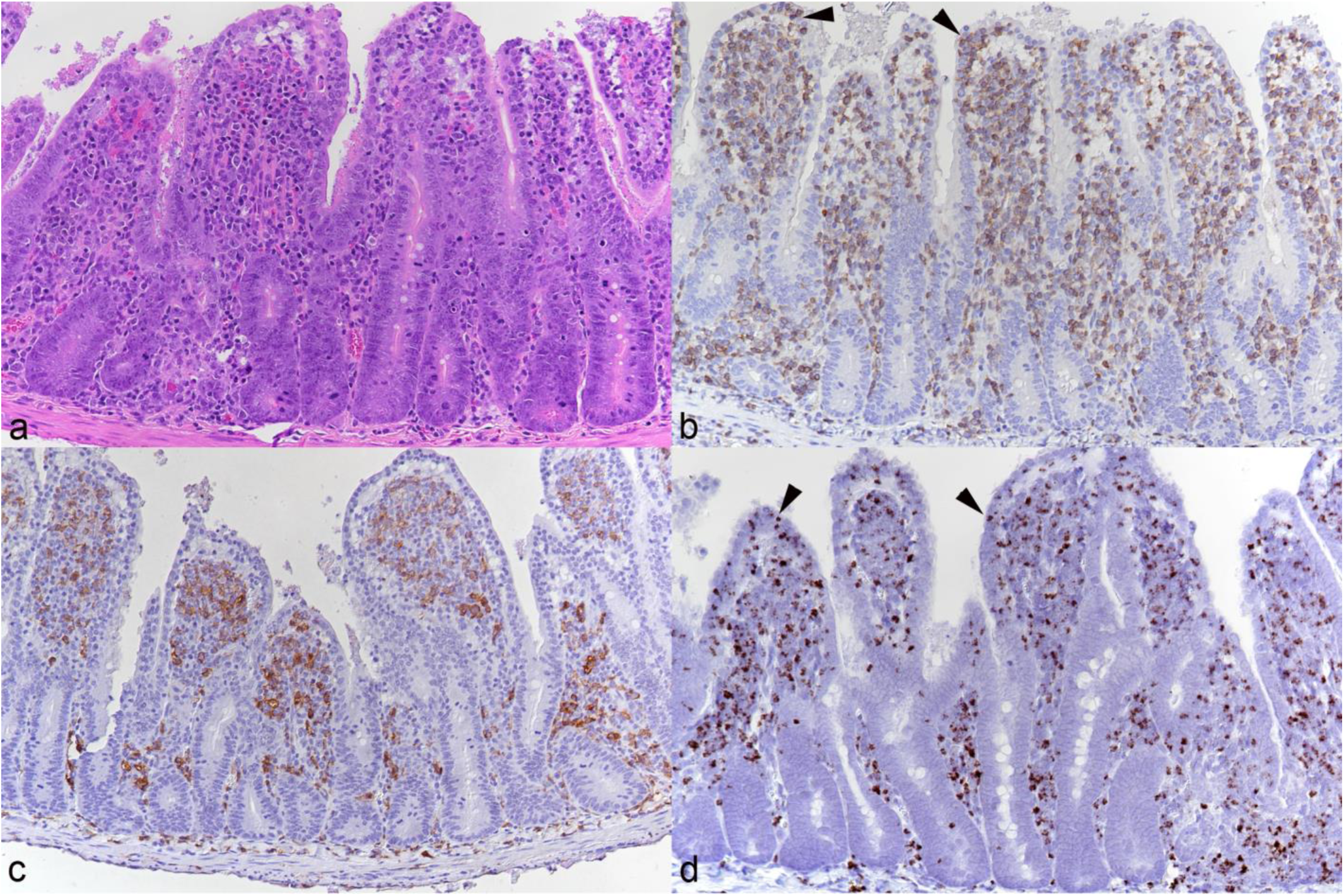
Small intestine, animal No. 3. **a.** Diffuse thickening of villi due to marked diffuse leukocyte infiltration of the mucosa. HE stain. **b.** The leukocyte infiltrate is dominated by T cells (CD3+) that are also infiltrating the epithelial layer (arrowheads). Immunohistochemistry (IH), hematoxylin counterstain. **c.** Macrophages (Iba1+) are the second most abundant leukocyte population in the mucosal infiltrate. IH, hematoxylin counterstain. **d.** Abundant infiltrating leukocytes, including some of those infiltrating the epithelial layer (arrowheads) are positive for viral RNA (Ov2.5). RNA-ISH, hematoxylin counterstain.

#### Nervous system

When the brain was examined (all animals from the main experiment; nos 5-10), a mild to moderate, focal extensive to diffuse mononuclear leptomeningitis was detected, with distinct perivascular arrangement of the infiltrating cells which were identified as T cells and macrophages. With more extensive meningeal infiltrates, involvement of the parenchyma (mononuclear encephalitis) was observed, represented by perivascular infiltrates of a similar composition (Fig. 7a-c), associated with microglial activation of the surrounding parenchyma which also contained individual migrated leukocytes, as staining for the T cell marker CD3 highlighted. A proportion of the infiltrating leukocytes carried viral RNA (Fig. 7d). With encephalitis, changes consistent with leukocytoclastic vasculitis were seen in all hamsters apart from animal no 9. Interestingly, the three available brains of the animals in the pilot study (animals no 2-4) only exhibited scattered T cells in the leptomeninges; however, some of these harbored viral RNA.

**Figure 7.**
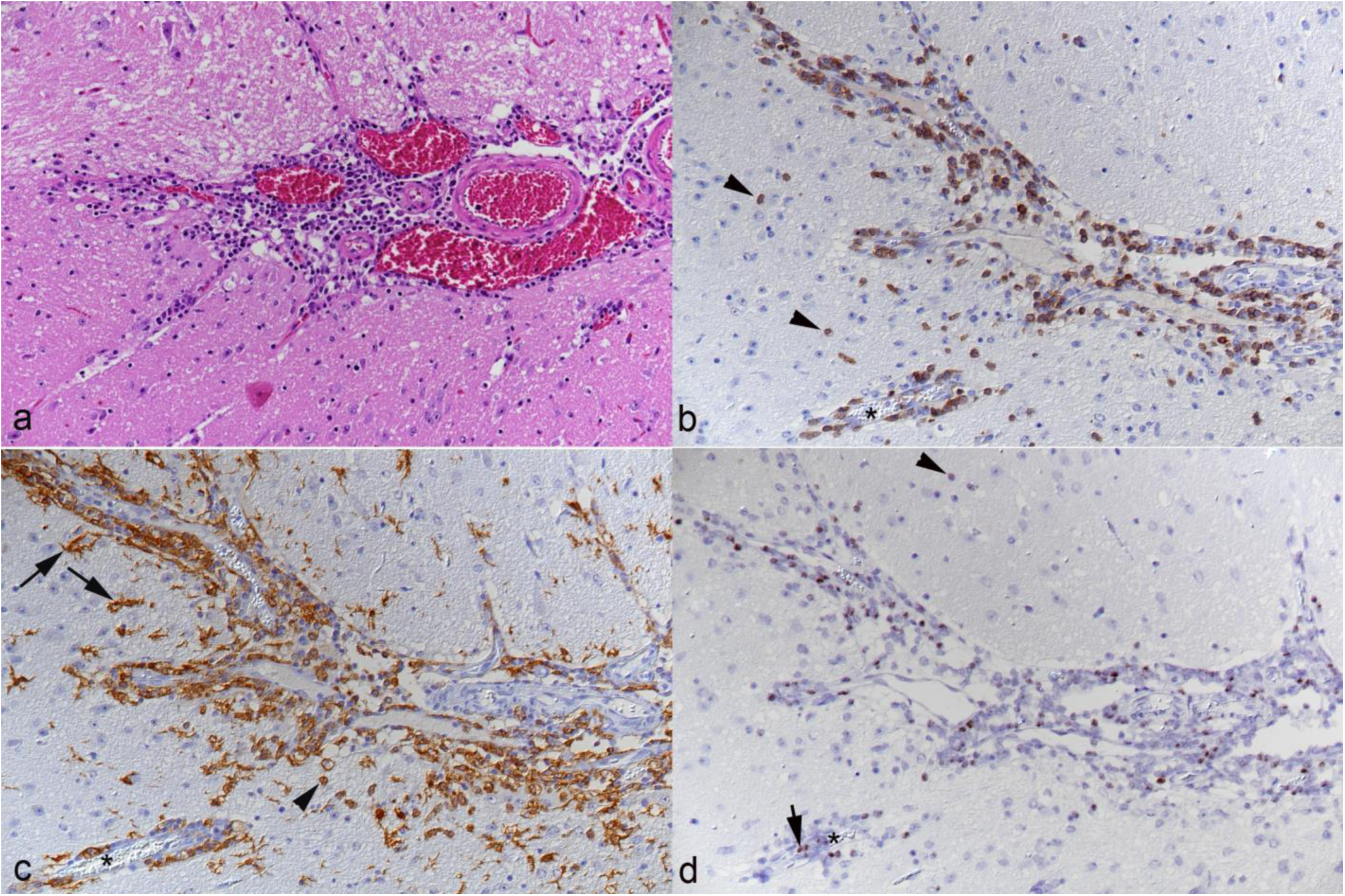
Brainstem, animal No. 10. **a.** Leptomeninx with moderate perivascular leukocyte infiltration. HE stain. **b.** Numerous leukocytes in the leptomeningeal and parenchymal perivascular infiltrates are T cells (CD3+). Some of these have migrated further in the parenchyma (arrowheads). Immunohistochemistry (IH), hematoxylin counterstain. **c.** Numerous leukocytes in the leptomeningeal and parenchymal perivascular infiltrates are macrophages (Iba1+). Some of these have migrated further in the parenchyma (arrowhead). The affected areas contain Iba1 positive cells with the morphology of activated microglia (arrows). IH, hematoxylin counterstain. **d.** A proportion of the leukocytes in the leptomeningeal and parenchymal perivascular infiltrates are positive for viral RNA (Ov2.5). Some of these have migrated further in the parenchyma (arrowhead). Arrow: Ov2.5 mRNA positive leukocyte in the blood. RNA-ISH, hematoxylin counterstain. Asterisks: affected parenchymal vessel.

In animals from the main experiment (nos 5-10), the <u>lumbar spinal cord</u>, a <u>sciatic nerve</u> and a *<u>M. biceps femoris</u>* were also examined. Mild leukomyelitis and leptomeningitis was consistently observed (Fig. 8). The latter was found to stretch to the spinal ganglia (periganglioneuritis) and was comprised of T cells and macrophages (Fig. 8b, c), with evidence of myelinophages (Fig. 8c), multifocal microgliosis (Supplemental Fig. S3) and mild satellitosis in the grey matter. Infiltrating leukocytes often carried viral RNA (Fig. 8d). In 5 of the 6 examined cases, the sciatic nerve exhibited mild focal perineurinal and occasionally perivascular neuronal T cell and macrophage infiltrates which contained cells positive for viral RNA (Supplemental Fig. S4). The biceps femoris muscle was generally unaltered; however, in three animals (nos 5, 7, 8) it exhibited very mild interstitial, perivascular leukocyte infiltrates.

**Figure 8.**
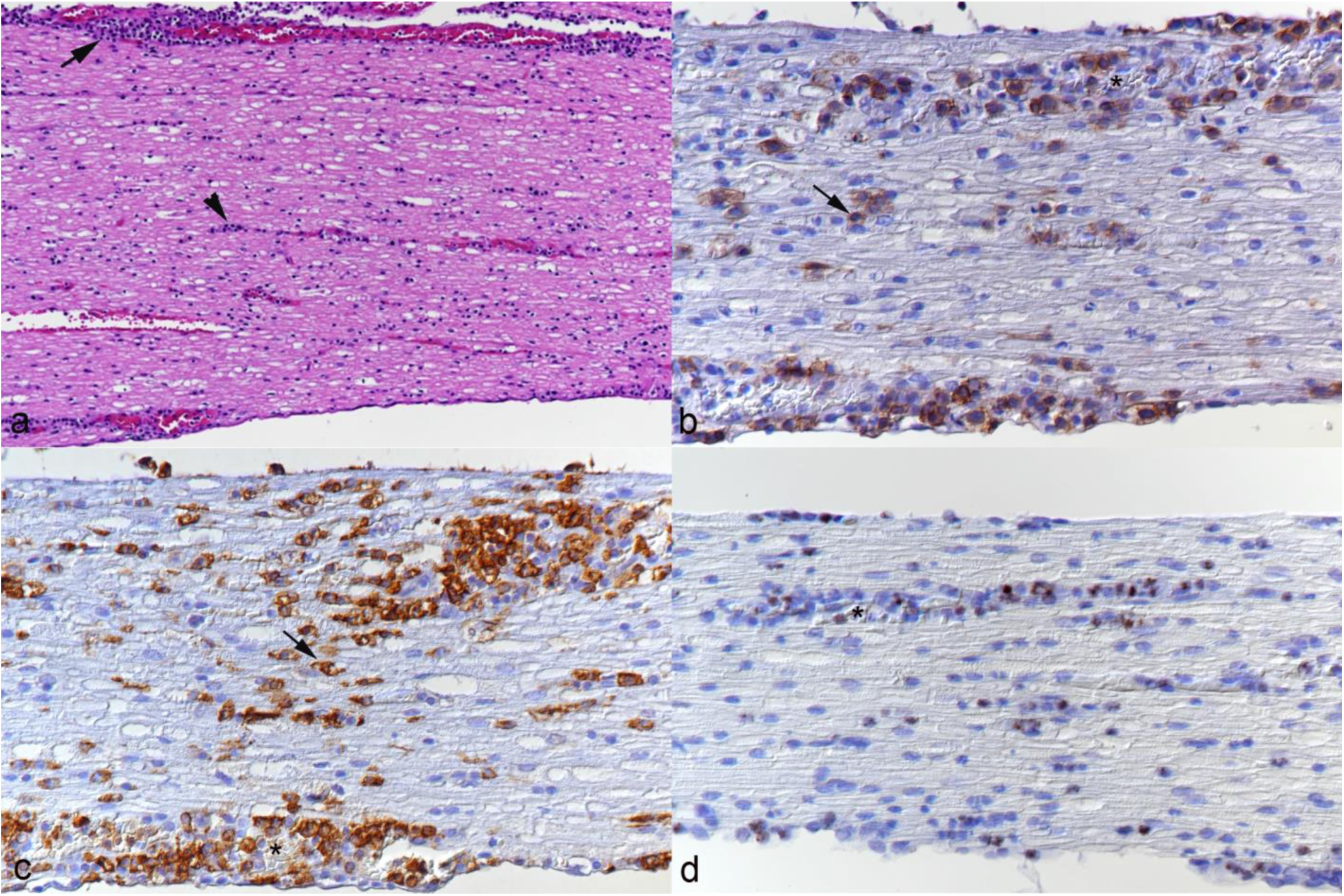
Lumbar spinal cord, longitudinal section, animal No. 10. **a.** Moderate multifocal leptomeningeal (arrow) and mild neuronal (arrowhead) perivascular leukocyte infiltration. HE stain. **b, c.** Area with intense inflammatory infiltration. **B.** T cells (CD3+) are abundant in the perivascular infiltrates, but are also present as individual cells between nerve fibers. Immunohistochemistry (IH), hematoxylin counterstain. **c.** Macrophages (Iba1+) are also abundant in the perivascular infiltrates, and there are Iba1 positive cells with the morphology of activated microglia (arrows) between nerve fibers. IH, hematoxylin counterstain. **d.** A proportion of the infiltrating leukocytes are positive for viral RNA (Ov2.5). RNA-ISH, hematoxylin counterstain. Asterisks: vessel lumen.

#### Kidneys (n=9)

In most animals (nos 2, 3, 5-10), the kidneys exhibited a mononuclear, T cell and macrophage composed, focal to multifocal interstitial infiltration and/or a subepithelial infiltration of the renal pelvis, consistent with a pyelitis. The infiltration varied in its extent from very mild to moderate and contained a variable proportion of leukocytes positive for viral RNA.

In animal No. 6, the urinary bladder was also examined histologically due to macroscopic changes. Histologically, a mild to moderate cystitis with mucosal and submucosal heterophilic and lymphohistiocytic infiltrates extending around the proximal part of the urethra and into the surrounding adipose tissue was seen.

#### Skin (n=3)

Skin from the dorsum was histologically examined in three animals from the pilot study (nos 2-4). It did not exhibit any histological changes; hence the skin was not examined in animals in the main experiment.

### OvHV-2 infects not only T cells and monocytes/macrophages but also selected squamous epithelial cells and possibly vascular endothelial cells

RNA-ISH, also in combination with immunohistochemistry for leukocyte markers, served to gain evidence of the target cell spectrum of OvHV-2 in the infected hamsters. This confirmed that circulating leukocytes carry viral RNA (Figs. 2d, 9) and hence spread the infection via viremia. It also highlighted that both T cells and monocytes in the blood are infected, evidenced by staining of cells in the liver sinuses (Fig. 9a, c). Both cell types appear to then transport the virus into the extravascular tissue, as shown in the portal infiltrates (Fig. 9b, d). In addition, there was also strong evidence of viral RNA in occasional vascular endothelial cells (Fig. 9e). Closer investigation of the squamous epithelium of the tongues and forestomachs also revealed the presence of viral RNA in scattered squamous epithelial cells in inflamed areas (Figs. 4d, 5f, 9f, g).

**Figure 9.**
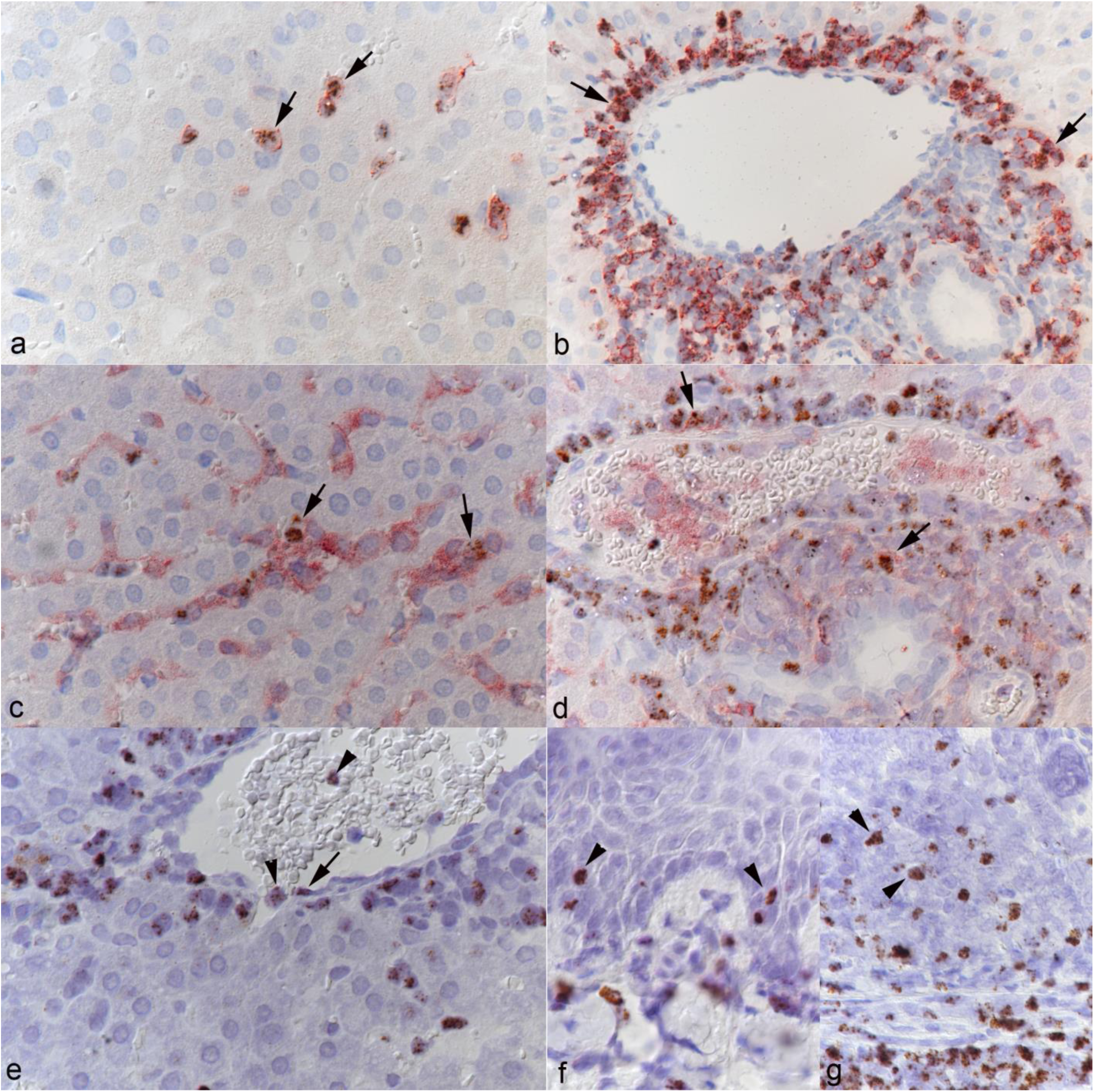
Target cells of OvHV-2 in the hamster. **a-d.** Liver, animal No. 10. **a.** There are several T cells (CD3+) in the sinuses that harbor OvHV-2 RNA (Ov2.5). **b.** Abundant T cells (CD3+) in the (perivenous) portal infiltrates harbor OvHV-2 RNA (Ov2.5). **c.** There are several monocytes (Iba1+) in the sinuses that harbor OvHV-2 RNA (Ov2.5). **d.** A few macrophages (Iba1+) in the (perivenous) portal infiltrates harbor OvHV-2 RNA (Ov2.5). **e.** Liver, animal No. 6. Central vein with leukocytes in lumen and adjacent sinus (arrowheads) positive for viral RNA (Ov2.5). There is also a cell with the morphology of an endothelial cell that harbors OvHV-2 RNA (arrow). **f, g.** Animal No. 1. **f.** Tongue. A few squamous epithelial cells exhibit a nuclear signal for OvHV-2 RNA (arrowheads). **g.** Forestomach. In addition to numerous infiltrating leukocytres, a few squamous epithelial cells exhibit a nuclear signal for OvHV-2 RNA (arrowheads). Immunohistochemistry and RNA-ISH, hematoxylin counterstain.

### Hemolymphatic tissue with OvHV-2 infection: evidence of expansion of the T cell compartments and the monocyte/macrophage pool

In all animals in the main experiment, the cervical, mediastinal and mesenterial <u>lymph nodes</u> were examined, from animals in the pilot experiment, individual lymph nodes were also included. All lymph nodes exhibited similar morphological features. Cortex and paracortex were generally poorly outlined due to a lack of distinct follicles and an expansion of the T cell compartment (Fig. 10a, b; Supplemental Fig. S5a, b).

**Figure 10.**
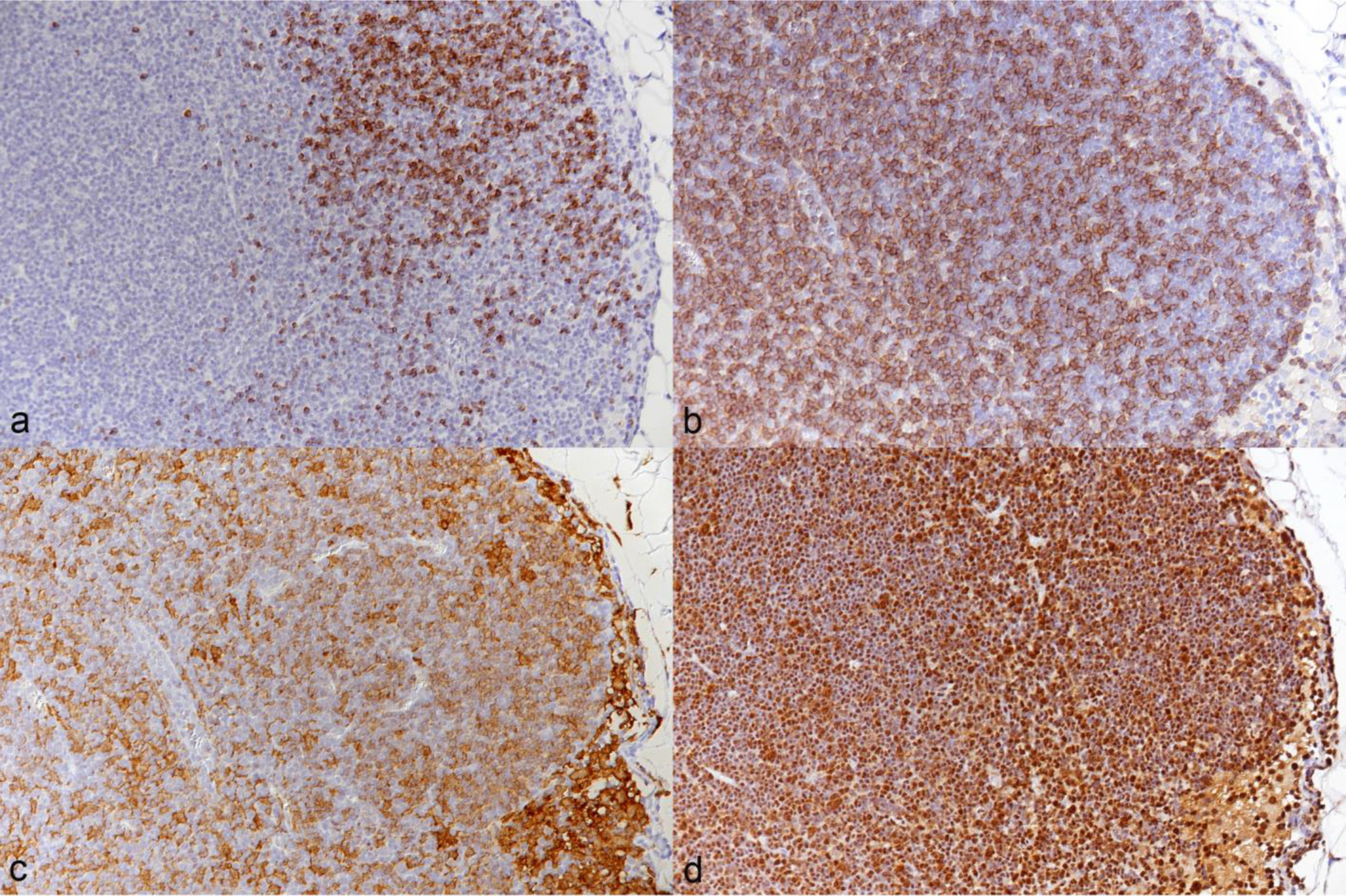
Mesenteric lymph node, animal No. 10. **a.** Staining for B cells (CD79a+) shows a lack of distinct follicles. Immunohistochemistry (IH), hematoxylin counterstain. **b.** Staining for T cells (CD3+) highlights expansion of the T cell compartment. IH, hematoxylin counterstain. **c.** Macrophages (Iba1+) expand the sinuses but are also abundant in areas populated by T cells and B cells. IH, hematoxylin counterstain. **d.** Staining for PCNA highlights abundant proliferating cells throughout the tissue. IH, hematoxylin counterstain.

Macrophages (Iba1+) were very abundant and appeared almost diffusely distributed (Fig. 10c; Supplemental Fig. S5c). Similarly, cell proliferation which was most pronounced in follicle centers in the control animals (Supplemental Fig. S5d) did not show any compartmental preference (Fig. 10d).

The <u>spleens</u> (n=9) exhibited a variable composition. In some animals (nos 2-5, 7), the white pulp was comprised of small primary follicles and small T cell zones, and the red pulp was of low cellularity. In the remaining animals (nos 6, 8-10), the follicles were of moderate size, and the T cell zones cell rich and poorly delineated (Fig. 11a, b; Supplemental Fig. S6a, b). The red pulp was of moderate to high cellularity which appeared to be due to an increase in monocytes/macrophages (Iba1+) and T cells (CD3+) among which were many proliferating (PCNA+) cells (Fig. 11b-d; Supplemental Fig. S6b-d).

**Figure 11.**
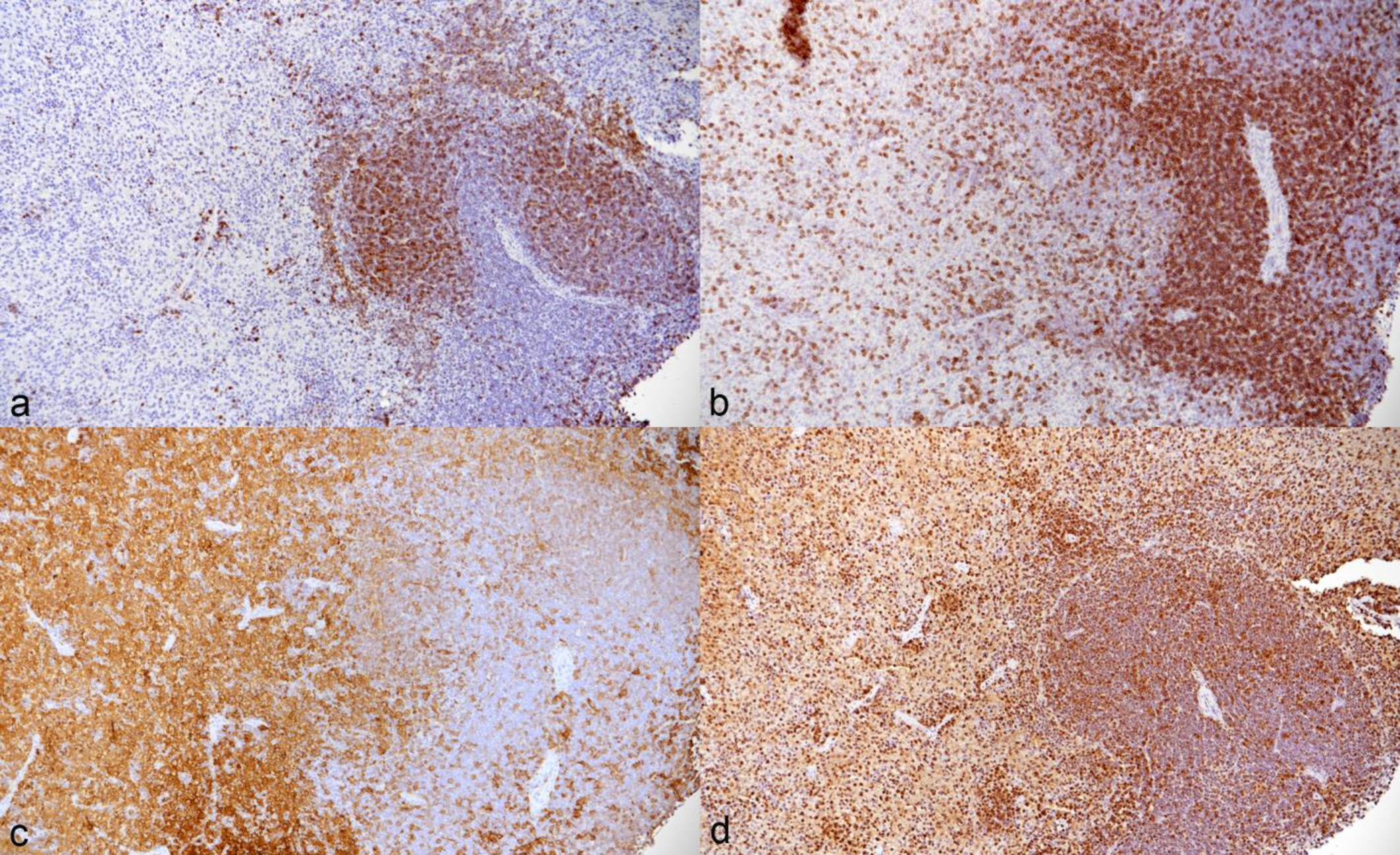
Spleen, animal No. 10. **a.** Staining for B cells (CD79a+) identifies small follicles. Immunohistochemistry (IH), hematoxylin counterstain. **b.** Staining for T cells (CD3+) highlights moderately sized T cell zones. T cells are also numerous in the red pulp. IH, hematoxylin counterstain. **c.** The red pulp contains very abundant monocytes/macrophages (Iba1+). IH, hematoxylin counterstain. **d.** Staining for PCNA highlights abundant proliferating cells in red and white pulp. IH, hematoxylin counterstain.

In both lymph nodes and spleen numerous disseminated leukocytes harbored viral Ov2.5 RNA (Fig. 12).

**Figure 12.**
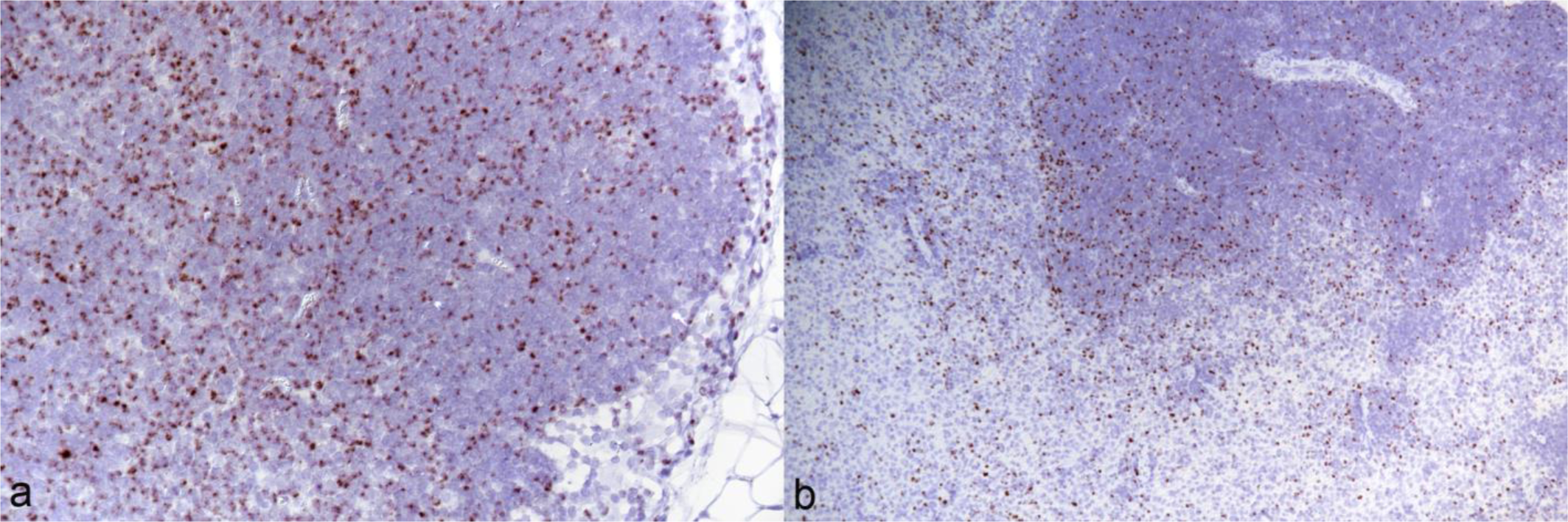
Lymphatic tissue, animal No. 10, RNA-ISH for Ov2.5, hematoxylin counterstain. **a.** Numerous disseminated leukocytes harbor viral RNA. **b.** Spleen. Numerous leukocytes in both red and white pulp harbor viral RNA.

Interestingly, whenever on the section, the adjacent mesentery exhibited mild or moderate perivascular mononuclear (T cells, macrophages) infiltrates with leukocytes that carried viral RNA.

The femoral and lumbar vertebral <u>bone marrow</u> (n=6; main study) was cell rich and appeared to contain more blastoid cells than in the age-matched control (Supplemental Fig. S7a, b). It comprised a substantial number of individual or small groups of T cells and numerous disseminated Iba1 positive cells, whereas in the control animal, T cells were basically absent and Iba1 positive cells were less abundant (Supplemental Fig. S6c-f).

## Discussion

MCF is a gammaherpesvirus induced sporadic disease of ruminants characterized by a variety of pathological changes affecting diverse organ systems but with a focus on blood vessels, skin and mucosa, and lymphatic tissues. The pathogenesis of these changes is not yet fully understood, indicating the need for suitable small animal models.

So far, the rabbit has mainly been promoted as an appropriate model of MCF. However, a search of the literature indicates that it might not recapitulate all relevant processes of natural MCF in cattle and other ruminants.^4,10,13,16,34,42,65^

The present study reports a renewed attempt at the hamster model of SA-MCF, making use also of *in situ* methodological approaches that were not available at the time when previous experimental infections were performed.^8^ It recapitulates and confirms the pathological effects of OvHV-2 in hamsters after intraperitoneal inoculation of lymphoid cells from infected animals but goes far beyond the previous studies as it provides further insight into the viral target cell spectrum and potential pathomechanisms underlying disease,^8^ also highlighting relevant pathogenic aspects that have so far not been considered to a major extent.

### Sustained elevated body temperature as the most reliable clinical sign

Persistent elevated body temperature has been used as a parameter to monitor the clinical disease after experimental infection.^10,30,57,76^ Indeed, it proved a reliable clinical sign of the disease in the hamsters also in the present study, similar to rabbits in previous studies,^34^ and was associated with viremia as determined by PCR on buffy coat cells from a first cohort of animals.

### T cell- and monocyte-mediated viremia and infection of hemolymphatic tissues

In the current study, hamsters were inoculated intraperitoneally with lymphoid cells isolated from spleen and lymph nodes of hamsters that had been experimentally infected with OvHV-2 in a prior experiment. Cells used in the initial study had been kept in liquid nitrogen for approximately 15 years; after thawing, live cells were selected and used for the inoculation at a dose of 2.2 x 10^5^ cells. As mentioned, in all inoculated animals elevation of the body temperatures indicated the onset of clinical disease.

Interestingly, the day of its onset post inoculation varied between the first and second experiment and between individual animals. Animals displayed onset of disease between 29-41 dpi and 15-17 dpi in the first and second experiment respectively, potentially due to the cells for the second experiment having been frozen down for a much shorter time before being used for infection.

Using RNA-ISH, our *in situ* examinations revealed that both circulating T cells and monocytes mediated the viremia. First evidence that OvHV-2 is spread systemically not only by T cells but also by monocytes was reported from experimental infections of sheep where some CD14+ monocytes were found to carry the virus.^45^ More recently, in our study on the arteritis in the rete mirabile of cattle with natural MCF, we found viral RNA in both circulating T cells and monocytes.^63^ The results of the present study not only confirm this finding but also detected numerous infected T cells and monocytes in lung capillaries and liver sinuses, indicating their abundance in the circulating blood. Further studies would be required to determine whether these findings are associated with hematological changes such as lympho- and monocytosis, in the infected hamsters.

Previous studies have suggested that in MCF the virus is carried by large granular lymphocytes (LGL) or large lymphoblasts, CD4+, CD8+ or CD4-/CD8-, that grow *in vitro* without exogenous cytokines and with an activated phenotype and consistent MHC-independent cytotoxicity;^7,10,64,71^ it was then not considered as clear whether these were T cells or NK cells.^71^ Similar to previous studies on naturally infected cattle with MCF,^54,63^ the present study cannot shed further light on the precise nature of the infected T cells as they likely all detected the fully assembled CD3 complex which would not be expressed by NK cells.^48^ However, taken together, their results suggest that T cells can be both lytically and latently infected.^54,63^

The literature frequently states that MCF is associated with hyperplasia of the lymphoid tissue, some papers have even called it a “lymphoproliferative disease”.^71^ However, some publications took a more differentiated approach and reported the changes in the lymphatic tissues of cattle and bison with MCF to range from hyperplasia to depletion; the former appears to mainly affect the T cell compartments.^22,51,76^ In both rabbits and hamsters, the most consistent reported morphological equivalent is an expansion of the T cell compartments in lymph nodes and spleen.^4,13,16,34,65^ This was confirmed in the present study, however, it also found evidence of expansion of the macrophage population and of increased proliferation of both (infected) T cells and macrophages in spleen and lymph nodes of OvHV-2 infected hamsters. Whether the release of infected cells from these tissues then provides the cells for the observed inflammatory processes remains to be clarified.

Our findings differ from the above described T cell restricted observations and are more in line with one experimental study in rabbits where increased proliferation was observed in all main cell types, T cells, B cells and monocytes, in spleen and mesenteric lymph nodes.^34^ In a previous experimental study in rabbits, a proportion of cells in the hyperplastic lymphoid tissue could be identified as CD8+, followed by lesser CD4+ cells. Since cells positive for either marker together made up less than the cells expressing a pan T cell marker, the authors concluded that, different from previous assumptions,^65^ the observed lymphoid hyperplasia was not due to expansion of the cytotoxic T cell pool but other, phenotypically altered lymphocytes.^4^ This hence represents another area of future research to clarify the pathogenesis of MCF.

The present study is the first to also examine the bone marrow of infected animals. Interestingly, it showed an increase in T cells (CD3+) and Iba1-positive cells (monocytes, possibly including also some immature stages) among the hematopoietic cells. This would be indicative of the presence of infected cells and, potentially, their proliferation which would suggest the bone marrow as a possible source of cell-associated viremia in animals with MCF.^2,5,6,73,75^ Unfortunately, we could not use our RNA-ISH approach on the bone marrow as the decalcification process destroys the RNA in the tissue (personal observation), however this is an area that warrants further investigation and could clarify whether the virus can persist in the bone marrow.

### Infected T cells and macrophages dominate the inflammatory infiltrates in MCF

The literature has provided variable information regarding the composition of the inflammatory infiltrates in both natural cases and animal models of MCF. An early light and electron microscopical study identified the vasculitis in MCF to be mediated by lymphocytes/lymphoblasts and fewer monocytes/macrophages.^40^ A subsequent study on two bovine MCF cases stated that macrophages made up more than half of the cells in the leukocyte infiltrates, accompanied by equal numbers or less T cells (CD8+, CD4+).^49^ A few years later, the investigation of each a calf and a bison with MCF found T cells (CD3+; predominantly CD8+, no CD4+ cells) to dominate the infiltrates, accompanied by low numbers of monocytes/macrophages.^68^

A dominance of T cells, with rare macrophages (CD14+) and B cells, has also been reported in the infiltrates in infected rabbits,^4^ and in MCF-associated inflammatory processes in the brain of cattle.^24^ However, our recent studies on cattle and a goat with MCF confirmed that, indeed, macrophages make up a large proportion of the (vascular) infiltrates.^42,63^ This is well reflected by the results obtained from the cohort of experimentally OvHV-2 infected hamsters, suggesting this species as highly suitable to study the development of the inflammatory processes.

### Vessel-centered inflammatory processes in MCF as a consequence of interaction between OvHV-2 infected leukocytes and endothelial cells?

Our recent study into the inflammatory processes in and around the rete mirabile arteries in cattle with SA-MCF has not only shown that it is driven by circulating infected T cells and monocytes, but also detected OvHV-2 infection in vascular endothelial cells and smooth muscle cells in the arterial media, suggesting that the virus plays a direct role in the recruitment of the inflammatory cells into the vascular wall and the perivascular compartment.^63^ A recent case report on SA-MCF in a goat provided further support for this interpretation and indicated that endothelial cells become more systemically infected, as it detected viral RNA (Ov2.5) and protein (OvHV-2 latency-associated nuclear antigen, oLANA) in lesional endothelial cells, and the latter also in endothelial cells of unaffected vessels.^42^ In the present study, we detected virus by RNA-ISH for Ov2.5 in endothelial cells in portal veins.

Like in natural SA-MCF in cattle and in the rabbit model,^13,16,34,65^ most inflammatory processes in infected hamsters were centered around blood vessels. They mainly represented perivascular infiltrates while vasculitis was observed but a less prominent feature than, e.g. the rete mirabile arteritis in cattle with MCF. Interestingly though, the hamsters consistently exhibited a valvular endocarditis affecting the left atrioventricular valves, mediated by (infected) T cells and monocytes. Taken together, the findings suggest that the inflammatory processes develop due to circulating activated, infected T cells and monocytes that home to tissues and emigrate from vessels prone to leukocyte emigration. Distribution of the lesions might either depend on actual endothelial cell infection or the susceptibility of vessels to endothelial activation and interaction with activated leukocytes. In this respect, MCF might be similar to Feline Infectious Peritonitis (FIP), a disease characterized by a phlebitis mediated and dominated by activated virus-infected monocytes. Interestingly, the phlebitis in FIP does not occur in all organs in affected cats but is rather stereotypically seen in leptomeninges, renal cortex (stellate veins) and eyes, and, less frequently, lungs and liver.^31^ The fact that with MCF, not only veins, where endothelial cells are known to better induce leukocyte adhesion than in arteries,^1,19^ are affected could indicate a more intense systemic endothelial cell activation in MCF than in FIP.^1^ The consistent affection of the atrioventricular valves in the hamsters in the present study is particularly interesting in this context. To our knowledge, virus-associated valvular endocarditis has so far only been reported with Coxsackie virus, an enterovirus, in individual patients, and in a mouse model of infection.^77^ Similarly, it is worth noting that the necrotic arteritis that is often seen in cattle,^40,51,76^ was not observed in our hamster cohort. It is also less marked in bison with MCF,^51,66^ and might, again like in FIP,^32^ depend on the extent of viremia and systemic inflammatory response in individual animals.

### Widespread inflammatory processes in the hamster model of SA-MCF

The present study included a detailed histological investigation of a wide range of tissues, taking into account the distribution of lesions reported in both natural MCF and animal models so far. As shown in Supplemental Table S1 which provides an overview of the reported distribution of (inflammatory) lesions in cattle with MCF and in experimentally OvHV-2 infected rabbits and hamsters, a rather wide range of organs is affected by perivascular and/or interstitial infiltrates, i.e. the liver (mainly portal areas), kidneys (interstitial infiltrates), lungs and heart with rather high consistency, whereas involvement of other organs is less frequent. In cattle, the brain is often involved, showing a nonsuppurative meningoencephalitis with or without lymphohistiocytic and, less often, necrotic vasculitis.^22,24^ We made similar observations in the hamster cohort, whereas in rabbits the brain and/or meninges seem not to be affected.^13^ In the present study, the histological examination of the spinal cord revealed a mild leukomyelitis and leptomeningitis, with periganglioneuritis of the spinal ganglia in all examined animals. The infiltrate had the same composition as elsewhere and included virus infected cells; it was associated with evidence of myelin degradation (presence of myelinophages), microgliosis and satellitosis. In most examined cases, the infiltrate also affected the sciatic nerve, and in some the biceps femoris muscle. Whether such inflammatory processes also occur with natural infections and in the rabbit is not known as the tissues have not been reported to be examined histologically. Interestingly, an older publication mentioned hind limb paralysis as a clinical sign in rabbits intravenously inoculated with cells carrying AlHV; however, a histological examination was apparently not performed.^30^ Taken together the findings indicate a more widespread distribution of the inflammatory processes than generally reported. However, neither the current study nor previous reports found evidence that OvHV-2 infects parenchymal cells in the brain or muscle cells.^24^

### Viral infection and inflammatory processes in the squamous epithelium

The present study laid a focus on the presence and type of changes in skin and the mucosa of the alimentary tract, taking into account that MCF is associated with rather consistent lesions at these sites in cattle and bison,^66,76^ and that we recently found erythema multiforme-like lesions in a goat with MCF.^42^ In cattle with MCF, the most detailed information on such lesions is available from an older study that investigated oral mucosa and esophagus in detail by light and electron microscopy. This characterized the (sub)epithelial infiltrates as mononuclear, dominated by lymphocytes/lymphoblasts, with fewer macrophages that increased in number with the severity of the lesions. Lymphocytes were found adjacent to epithelial cells and there was evidence of phagocytosis of degenerate epithelial cells. The authors also described acantholysis and necrosis of epithelial cells.^8,40,50^ The recently reported case of erythema multiforme like skin lesions in a goat with SA-MCF described ulceration, transepidermal apoptosis with satellitosis, interface dermatitis, folliculitis and dermal arteritis and found viral RNA (Ov2.5) not only in the nucleus of infiltrating leukocytes and vascular endothelial cells but also in keratinocytes, suggesting that a cell-mediated immune response against the virus might at least contribute to the development of the lesions.^42^ The present study consistently observed similar, partly ulcerative lesions on tongue and forestomachs of the infected hamsters, comparable to an older study.^8^ Like the skin of the goat with natural SA-MCF, the affected squamous epithelium exhibited epithelial cells dying by apoptosis, as confirmed by the expression of the apoptosis marker cleaved caspase-3.^42^ It is not unlikely that the cell death described in the earlier paper does indeed represent apoptosis.^40^ Indeed, a review paper on the pathology of SA-MCF has shown images from the urinary bladder and oral mucosa of an affected bison, with apoptosis of epithelial cells.^51^ The older detailed study also showed that epithelial lesions are widespread in cattle and affect the entire gastrointestinal tract, the conjunctiva, the respiratory tract, various glandular ducts, the urinary bladder and the choroid plexus, with degenerative epithelial changes exclusively at sites of inflammatory infiltrations.^40,41^ Despite the presence of vasculitis and/or perivascular infiltrates in the deeper lamina propria and the earlier assumption that the lesions could result from vascular disruption,^62^ the authors concluded that due to the lack of substantial infarcts and vascular thrombi, ischemia does likely not play a major role in the pathogenesis of the epithelial lesions. Instead, they saw substantial similarity to lesions seen in graft-versus-host disease.^40^ The results of the present study and the composition of the inflammatory infiltrates support this interpretation, considering also that a previous report mentioned a dominance of macrophages in vascular and epithelial lesions in two natural cases of MCF in cattle.^49^ Interestingly, while we detected apoptotic cells in lesional epidermis/squamous epithelium in both the goat with MCF and our hamsters, we did not find morphological evidence that latently infected epithelial cells underwent apoptosis.^42^ This is in line with our recent findings in MCF vasculitis in cattle,^63^ and another study that did not even report damage in lytically infected cells.^54^ This provides further evidence that any virus induced changes, whether functional or phenotypic, do not present as direct cytopathic effects. Whether the epithelial cells were latently infected or even promote lytic infection could not be determined, as Ov2.5, the target of our RNA-ISH encoding a protein similar to ovine interleukin 10, is expressed during both viral phases.^3,29^ It is also not clear which role the infection of the infiltrating leukocytes might play in this scenario. Does it require infected, activated T cells and monocytes/macrophages to home to the tissue (i.e. squamous epithelium) and carry infection to epithelial cells, or is it the infection of the epithelium that attracts the leukocytes? Does the (accidental) recruitment of virus infected T cells and monocytes through the vasculitis foster or potentiate the mucosal/epidermal disease processes?

Interestingly though, while we observed a strong diffuse mucosal infiltration of the intestine by virus infected T cells and macrophages, with villus blunting and fusion, we did not find evidence of viral infection of enterocytes. Similarly, we did not detect viral RNA in respiratory or alveolar epithelial cells which suggests a selective epithelial tropism of the virus at least in hamsters. It remains open whether the strong infiltration of the intestinal mucosa occurs because this compartment contains abundant leukocytes and in particular residential T cells anyway, rendering it prone to more intense leukocyte recruitment. In experimentally (BJ 1035 cells) infected rabbits, RNA-ISH provided evidence of viral replication in epithelial cells in the appendix.^46^ In this study, OvHV-2 ORF63 mRNA was detected, and ORF-63 is predicted to code for a viral tegument protein.^46^ However, since this study did not detect any other cells with an ORF63 mRNA signal in any of the other examined tissues (mesenteric lymph node, lung, spleen, liver, kidney) it is difficult to compare with other studies, including ours.^54^

Interestingly, using a mouse monoclonal antibody (15-A) against a conserved epitope of MCF viruses, binding a 45 kDa protein of a larger glycoprotein complex,^38^ a previous study on cattle with SA-MCF detected OvHV-2 antigen in the cytoplasm of a wide range of epithelial cells (renal tubules, degenerating bile ducts, small intestinal degenerating crypts), in cells with macrophage morphology in the mucosa of the small intestine with inflammatory infiltration and in the mesenteric lymph node, and in endothelial cells in renal medullary capillaries but not in the brains despite the presence of meningoencephalitis and vasculitis.^21^ The epithelium of the respiratory tract was not found to be positive, despite its consistent infection in the acute phase of most gammaherpesviruses in their natural hosts.^25,74^

The present study cannot yet address the key question regarding the pathogenesis of MCF, as it did not investigate the trigger for the proliferation and activation of infected leukocytes and hence the inflammatory processes. It can also not provide answers to which extent the virus targets vascular endothelial cells and (squamous) epithelia and whether the recently suggested even broader target cell spectrum is of major relevance for the development of disease.^21^ However, it offers the hamster as a suitable experimental host for studies on the pathogenesis of SA-MCF; with its help it should be possible to further investigate the interplay between virus and host cells and between infected cells, with particular emphasis on a potential graft-versus-host reaction.

## Acknowledgements

We are grateful to the technical staff in the Histology Laboratory, Institute of Veterinary Pathology, Vetsuisse Faculty, University of Zurich (IVPZ), for excellent technical support. We would also like to thank Dr Paula Grest and Dr Frauke Seehusen, IVPZ, for fruitful discussions.

