## Supplemental Table S1 for "The golden Syrian hamster (*Mesocricetus auratus*) as a model to decipher relevant pathogenic aspects of sheep-associated malignant catarrhal fever"

**Supplemental Table S1**. Pathological changes reported in systematic studies in natural SA-MCF in cattle or summarized in a recent textbook,[16] in the rabbit model of SA-MCF, and in the hamster model as observed in the present study, complemented by findings reported in a previous publication.[2]

| **Organ/tissue** | **Natural MCF (cattle)** | **Rabbit model** | **Hamster model** |
|---|---|---|---|
| Vessels and heart | Vessels: disseminated mononuclear vasculitis and pv accumulations[8,12,15,16]<br>Heart: lymphoplasmacytic myocarditis[8] | Vessels: general arteriolitis-phlebitis in bronchial circulation[4,10]<br>Heart: occasional interstitial LC accumulations[6] | Vessels: vasculitis in occasional portal veins (liver) and of vessels in brain parenchyma with encephalitis, widespread pv infiltrations (TM[b])[c]<br>Heart: variable degree of endocarditis and myocarditis[2]; valvular endocarditis (AV valves), occasional parietal endocarditis and myocardial interstitial infiltrates (TM[b])[c] |
| Alimentary tract mucosa | Erosive/ulcerative lesions and/or lymphoplasmacytic inflammatory infiltrates: tongue, gingiva, oral mucosa, esophagus, forestomachs, abomasum, intestines[8,11,13,15] | Esophagus: focal subepithelial mononuclear lymphoid cell accumulations[a] associated with ballooning degeneration of the epithelium[14]<br>Intestines: information limited to appendix in areas of lymphoid structures (see below) | Oral mucosa: inflammatory processes tongue, cheek pouches, hard palate[2]; subepithelial infiltrates in tongue[d] (TM[b])[c]<br>Esophagus: subepithelial infiltrates (TM[b])[c]<br>Non-glandular forestomach: acantholysis and hyperkeratosis[2]; erosive/ulcerative proventriculitis[d], marked submucosal infiltration (TM[b])[c]<br>Stomach: mild/moderate infiltration of the mucosa (TM[b])[c;2]<br>Small intestine: marked diffuse mucosal infiltration (TM [b])[c;2]<br>Caecum: ulcerative lesions[2] |
| Liver | Pv mononuclear infiltration in portal areas[8,16] | Peribiliary and perivascular lymphoid cell accumulations[a14]; periportal lymphoid cell accumulations (mainly T cells)[1,3]; portal hepatitis (large LC, fewer macrophages, plasma cells and heterophils), arteritis and phlebitis, biliary hyperplasia, focal areas of hepatocellular necrosis[6]; (peri)cholangitis, portal phlebitis-arteritis, hepatic necrosis, granulomatous lesions[10] | Moderate portal infiltrates (TM[c])[2] |
| Lymphatic tissue, bone marrow | Lymph nodes: variable; hyperplasia in T cell dependent areas in interfollicular | Lymph nodes (MLN): expansion of the paracortex[14]/interfollicular zones (T cells)[3]; marked hyperplasia of cortex, paracortex and lymphoid | Lymph nodes: lymphadenitis and hyperplasia[2]; expansion of T cell compartment and macrophage population, with proliferation of cells[c] |

| | | | |
|---|---|---|---|
| | cortical and paracortical zones, depletion[8,16]<br>Spleen: variable, ranging from lymphoid cell hyperplasia in PALS to depletion[8,16] | follicles, focal necrosis (mainly T cells)[1]; mild expansion of paracortex (and cortex) by large LC[6]; diffuse lymphadenitis with progression to granulomatous lymphadenitis[10]<br>Lymphoid tissue (appendix): hyperplasia of T cell areas, necrosis[14]; marked hyperplasia of lymphoid follicles and interfollicular areas (mainly T cells), necrosis of lymphoid follicles[1]<br>Spleen: hyperplasia of PALS and red pulp (mainly T cells)[1,3]; mild increase in large LC in PALS[6] | Spleen: prominent PALS[2]; variable expansion of T cell zones, increased cellularity of red pulp (TM[b])[c]<br>Bone marrow: cell rich; with substantial numbers of T cells and numerous monocytes (Iba1 positive cells)[c]<br>Thymus: premature involution[2] |
| Central and peripheral nervous system, eyes | Brain: nonsuppurative meningoencephalitis with(out) lymphohistiocytic vasculitis and/or necrotic vasculitis[8,9]<br>Spinal cord, peripheral nerves: no lesions described/not examined<br>Eyes: lympho(plasma)cytic conjunctivitis, keratitis, iridocyclitis, uveitis[8,12] | Brain: no evidence of meningitis and/or encephalitis[4]<br>Spinal cord, peripheral nerves: not examined<br>Eyes: lymphoid hyperplasia in palbebral conjunctiva;[3] gross evidence of keratoconjunctivitis;[10] no evidence of keratoconjunctivitis[4] | Brain: mononuclear leptomeningitis and encephalitis (TM[c]); mild lymphoid meningitis[2]<br>Spinal cord: lymphohistiocytic leukomyelitis, leptomeningitis and periganglionitis (TM[b])[c]<br>Sciativ nerve: mild focal neuritis and perineuritis (TM[b])[c]<br>Eyes: mononuclear infiltration of conjunctival lamina propria with acanthosis and hyperkeratosis at conjunctival-epidermal junction[2] |
| Respiratory tract | Nose: (ulcerative) lymphoplasmacytic rhinitis[7,8]<br>Airways: erosive tracheobronchitis[16]<br>Lungs: interstitial pneumonia, bronchopneumonia[8] | Nose: no lesions described (not examined)<br>Trachea: lymphoid cell accumulations[a14]<br>Lungs: perivascular and peribronchial/-bronchiolar lymphoid cell accumulations (mainly T cells);[1] interstitial pneumonia with intralesional phlebitis-arteritis[10] | Nose: not examined<br>Trachea: focal mononuclear infiltrates[2]<br>Lungs: arteritis and focal interstitial lymphoid infiltrates[2]; increase in TM[b] in capillaries, some leukocyte emigration and perivascular accumulation[c] |
| Urinary tract | Kidneys: pv mononuclear infiltrations in cortex, lymphocytic interstitial nephritis;[8,16]<br>Urinary bladder: (ulcerative) lymphoplasmacytic or haemorrhagic cystitis[8,11] | Kidneys: lymphoid cell accumulations[a];[14] frequent perivascular LC accumulations in cortex (mainly T cells)[1]; occasional interstitial LC accumulations and occasional vasculitis.[6]<br>Urinary bladder: pv LC accumulations[6] | Kidney: focal interstitial infiltrations and/or focal pyelitis (TM[b]) [c]<br>Urinary bladder: ulcerative cystitis;[2] mild to moderate heterophilic cystitis[c] |
| Skin | Ulcerative/necrotic dermatitis in several locations; lichenoid infiltration in upper dermis, stretching into epidermis[5,16] | Not examined | Occasional mononuclear infiltrates in dermis, degeneration of basal cells and hyperkeratosis of overlying epidermis[2]; no changes observed[c] |
| Other locations | Variable degree of mononuclear infiltration beneath and between epithelial cells in respiratory tract, entire gastrointestinal tract, biliary epithelium, | Occasional interstitial LC accumulations in adrenal glands, thyroid glands, pancreas, tongue, trachea, sclera[6] | Skeletal muscles: occasional necrosis and interstitial mononuclear infiltration (diaphragm and cremaster)[2]; very mild interstitial, pv leukocyte infiltrates[c] |

| | |
|---|---|
| | glandular ducts, choroid plexus[11]; lymphoplasmacytic sinusitis, thyroiditis[8] |

Legend: LC – lymphocyte(s); PALS – periarteriolar lymphoid sheaths; pv - perivascular

[a]The lymphoid cells in non-lymphoid tissues were CD43+ T cells, as determined by immunohistochemistry.
[b]TM: The infiltrate is comprised of T cells (CD3+) and macrophages (Iba1+) of which a proportion is infected, as shown by RNA-ISH for Ov2.5.
[c]Results obtained in the present study
[d]The infiltrates are accompanied by apoptosis of epithelial cells.
