## Supplemental Table S2 for "The golden Syrian hamster (*Mesocricetus auratus*) as a model to decipher relevant pathogenic aspects of sheep-associated malignant catarrhal fever"

**Supplemental Table S2**. Antibodies, antigen retrieval and detection methods used in immunohistology

| Antigen | Antibody (clone) | Dilution (incubation) | Pretreatment | Detection method |
|---|---|---|---|---|
| CD3 | Rabbit mAb (SP7)[a] | 1: 400 (1 h, 37 °C) | EDTA[b] | Discovery Chromo Map (OmniMAP anti-Rabbit)[b] |
| CD79a | Mouse mAb (MH57)[c] | 1:1,000 (1h, RT) | EDTA | EnVision Mouse[d] |
| Iba-1 | Rabbit pAb[e] | 1:1,000 (1h, RT) | Citrate | EnVision Rabbit[d] |
| PCNA | Mouse mAb (PC10)[d] | 1:800 (ON, 4 °C) | Citrate | MACH-4[f] |
| Cleaved caspase 3 | Rabbit mAb[g] | 1:200 (ON, 4 °C) | Citrate | EnVision Rabbit[d] |

Legend: Iba-1, ionized calcium binding adaptor molecule 1; pAb, polyclonal antibody; mAb, monoclonal antibody; ON – overnight; PCNA – proliferating cell nuclear antigen; RT – room temperature.

EDTA: 20 min incubation in EDTA buffer (pH9; Dako/Agilent) in a pressure cooker at 98 °C.

Citrate: 20 min incubation in citrate buffer (pH 6; Dako/Agilent) in a pressure cooker at 98 °C.

[a] Bioscience
[b] Ventana/Roche
[c] Bio-Rad
[d] Dako/Agilent
[e] Wako
[f] Biocare Medical
[g] Cell Signaling Technology
.
