## Supplemental Table S3 for "The golden Syrian hamster (*Mesocricetus auratus*) as a model to decipher relevant pathogenic aspects of sheep-associated malignant catarrhal fever"

**Supplemental Table S3**. Pilot study, information on the day of euthanasia post challenge, body temperature at euthanasia, and organs affected by the histologically observed leukocyte infiltrates. The histological changes are described in the main manuscript; here only those organs are listed that showed leukocyte infiltrates (i.e. T cell and macrophage infiltrates with viral mRNA signal).

| Animal | Euthanasia (dpi) | Body temperature | Affected organs |
|---|---|---|---|
| 1 | 29 | 37.4 °C | Lungs |
| 2 | 33 | 37.4 °C | Tongue, stomach, SI, liver, lungs, kidneys, brain |
| 3 | 41 | 37.8 °C | Tongue, stomach, SI, liver, lungs, kidneys, brain |
| 4 | 41 | 36.2 °C | Tongue, stomach, SI, liver, lungs, brain |
| 5 | 17 | 31.7 °C | Tongue, stomach, SI, liver, heart, lungs, kidneys, brain, SC |
| 6 | 17 | 38.0 °C | Tongue, stomach, SI, liver, heart, kidneys, brain, SC, SN |
| 7 | 17 | 38.0 °C | Tongue, stomach, SI, liver, heart, kidneys, brain, SC, SN |
| 8 | 15 | 37.5 °C | Tongue, stomach, SI, liver, heart, kidneys, brain, SC, SN |
| 9 | 17 | 37.7 °C | Tongue, stomach, SI, liver, heart, kidneys, brain, SC, SN |
| 10 | 17 | 37.8 °C | Tongue, stomach, SI, liver, heart, kidneys, brain, SC, SN |

**Initial study** (animals 1-4): From animal 1, only the lungs were examined histologically, from animals 2-4, skin (dorsum), tongue, stomach, small intestine (SI), liver, spleen, lungs, kidneys, and the brain was examined.

**Second study** (animals 5-10): From all animals, tongue, oesophagus, stomach, small intestine (SI), liver, heart, trachea, lungs, kidneys, brain, lumbar spinal cord (SC), right sciatic nerve (SN) and *Musculus biceps brachii* as well as the spleen and the cervical, mediastinal and mesenteric lymph nodes were examined.
