## Supplemental Figures for "The golden Syrian hamster (*Mesocricetus auratus*) as a model to decipher relevant pathogenic aspects of sheep-associated malignant catarrhal fever"

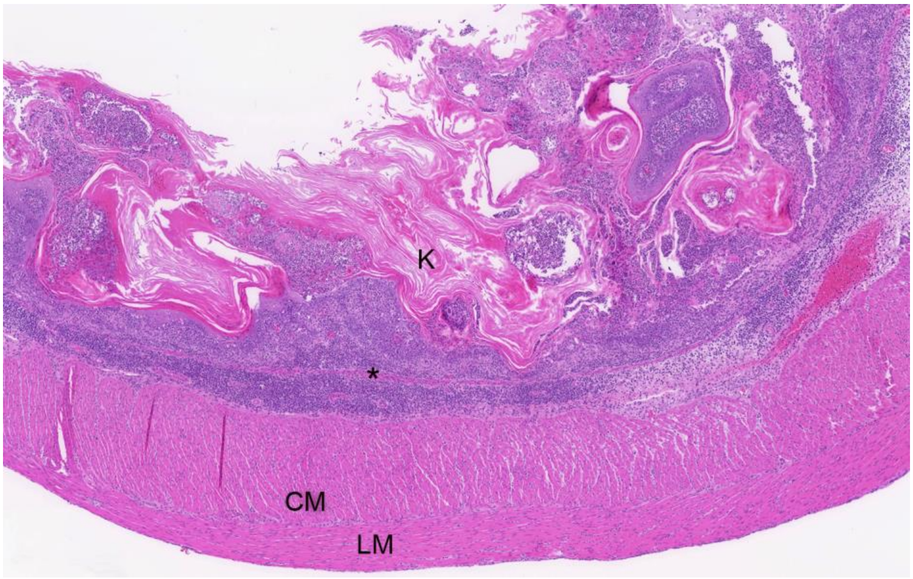


**Figure S1**. Forestomach, animal No 3. Overview of an area with extensive serocellular crust formation and marked mucosal and submucosal mononuclear infiltration. K: keratin layer; CM: circular muscle layer; LM longitudinal muscle layer; asterisk: muscularis mucosae.

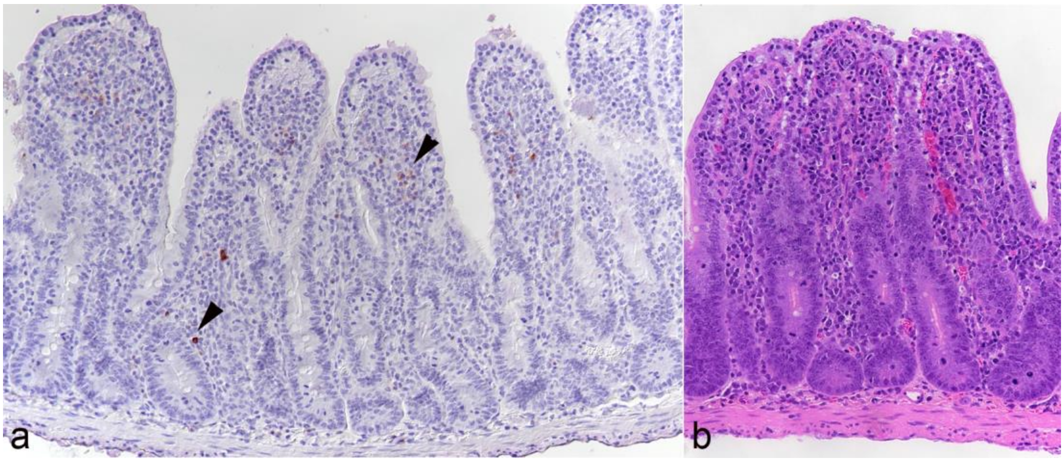


**Figure S2**. Small intestine, animal No 3. **A.** Among the infiltrating leukocytes in the mucosa are scattered CD79a positive B cells and plasma cells (arrowheads) Immunohistochemistry (IH), hematoxylin counterstain. **B.** The intense mucosal infiltration is associated with bluting and fusion of the thickened villi. HE stain.

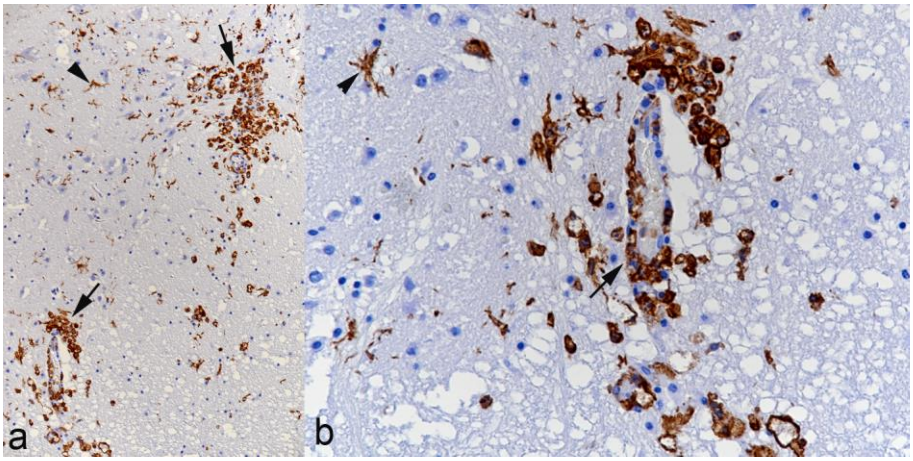


**Figure S3**. Lumbar spinal cord, animal No. 7. Immunohistochemistry for Iba1, hematoxylin counterstain. **a.** There are focal areas of microgliosis (arrows) and activated microglial cells (arrowhead) in the parenchyma. **b.** A closer view of a vessel with surrounding parenchyma confirms monocyte emigration and perivascular macrophage infiltration (arrow) and the presence of activated microglial cells (arrowhead).

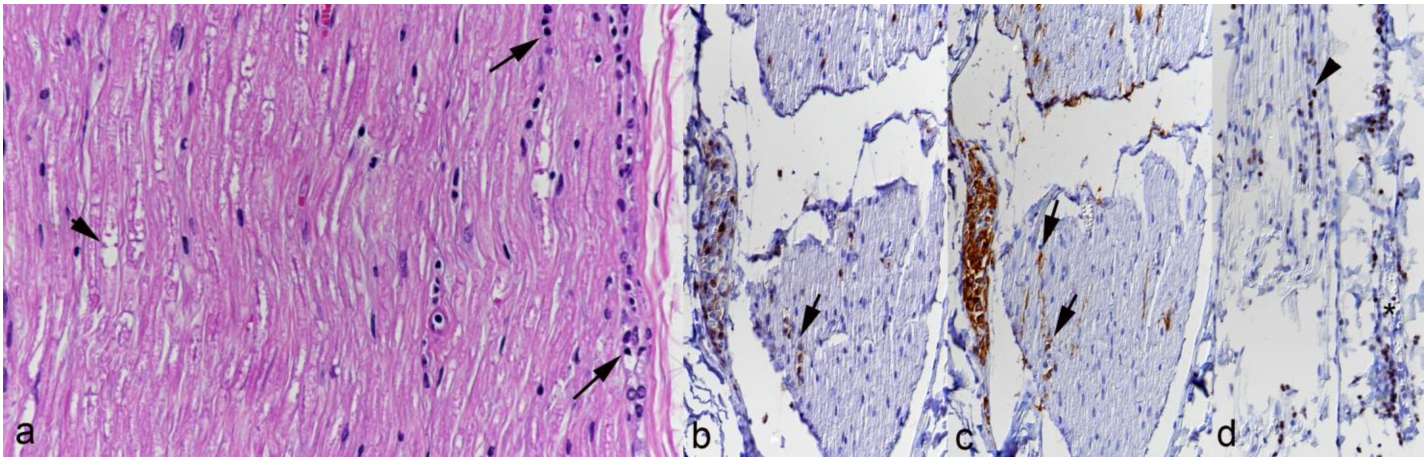


**Figure S4**. Sciatic nerve, animal No. 10. **a.** There are individual degenerate nerve fibers (arrowhead) and a few infiltrating leukocytes (arrows). HE stain. **b.** T cells (CD3+) infiltrating between nerve fibers (arrow). Immunohistochemistry (IH), hematoxylin counterstain. **c.** Macrophages (Iba1+) infiltrating between nerve fibers. IH, hematoxylin counterstain. **d.** Infiltrating leukocytes harbor OvHV-2 RNA (Ov2.5). RNA-ISH, hematoxylin counterstain.

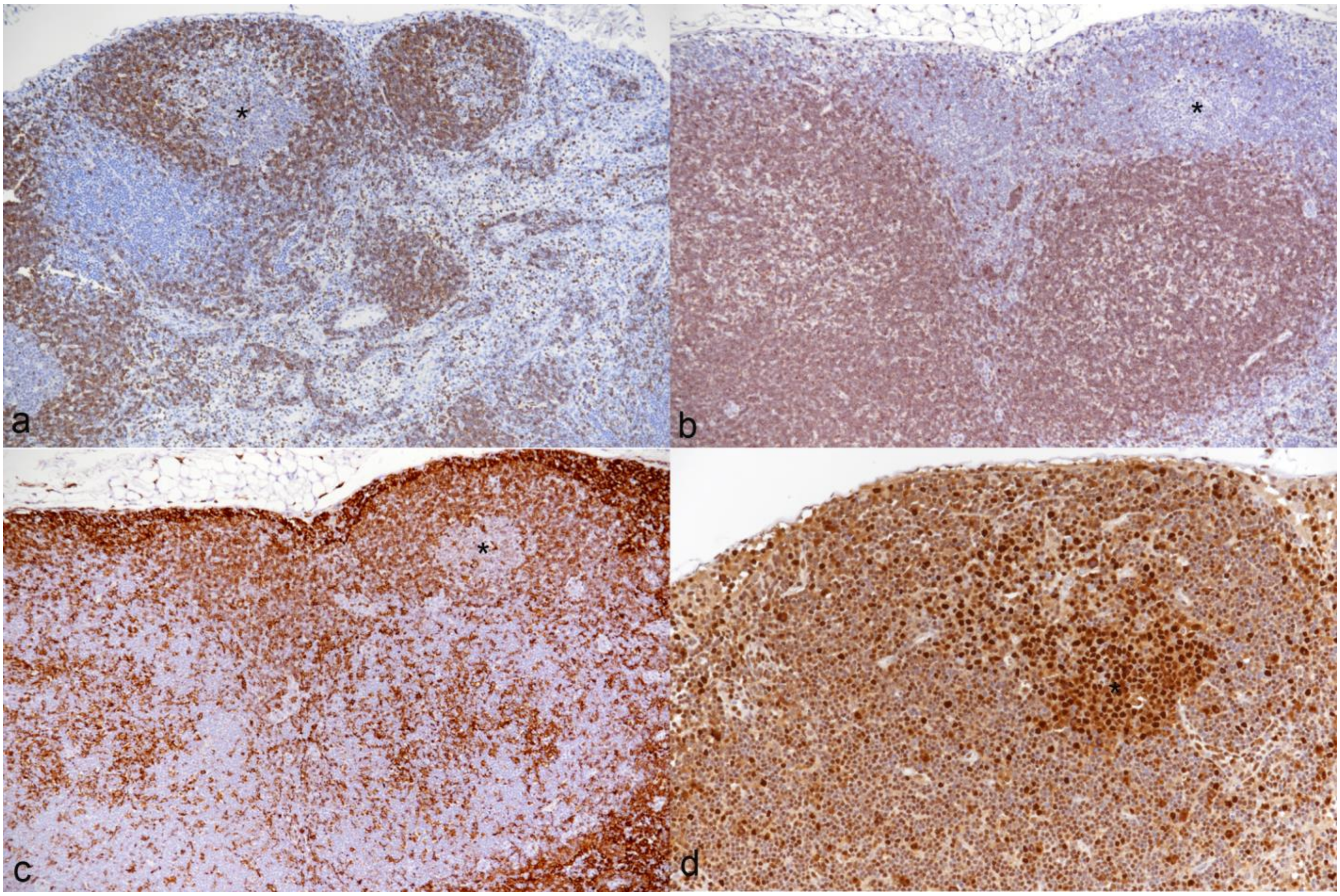


**Figure S5**. Mesenteric lymph node, control animal. The asterisk highlights a follicular germinal center. **a.** Staining for B cells (CD79a+) shows several distinct follicles comprising the cortex. Immunohistochemistry (IH), hematoxylin counterstain. **b.** Staining for T cells (CD3+) highlights a well defined paracortex (T cell compartment). IH, hematoxylin counterstain. **c.** Macrophages (Iba1+) are mainly seen in sinuses and in the cortex, they are less numerous in the paracortex. IH, hematoxylin counterstain. **d.** Staining for PCNA highlights pronounced proliferation in the germinal center of a follicle. IH, hematoxylin counterstain.

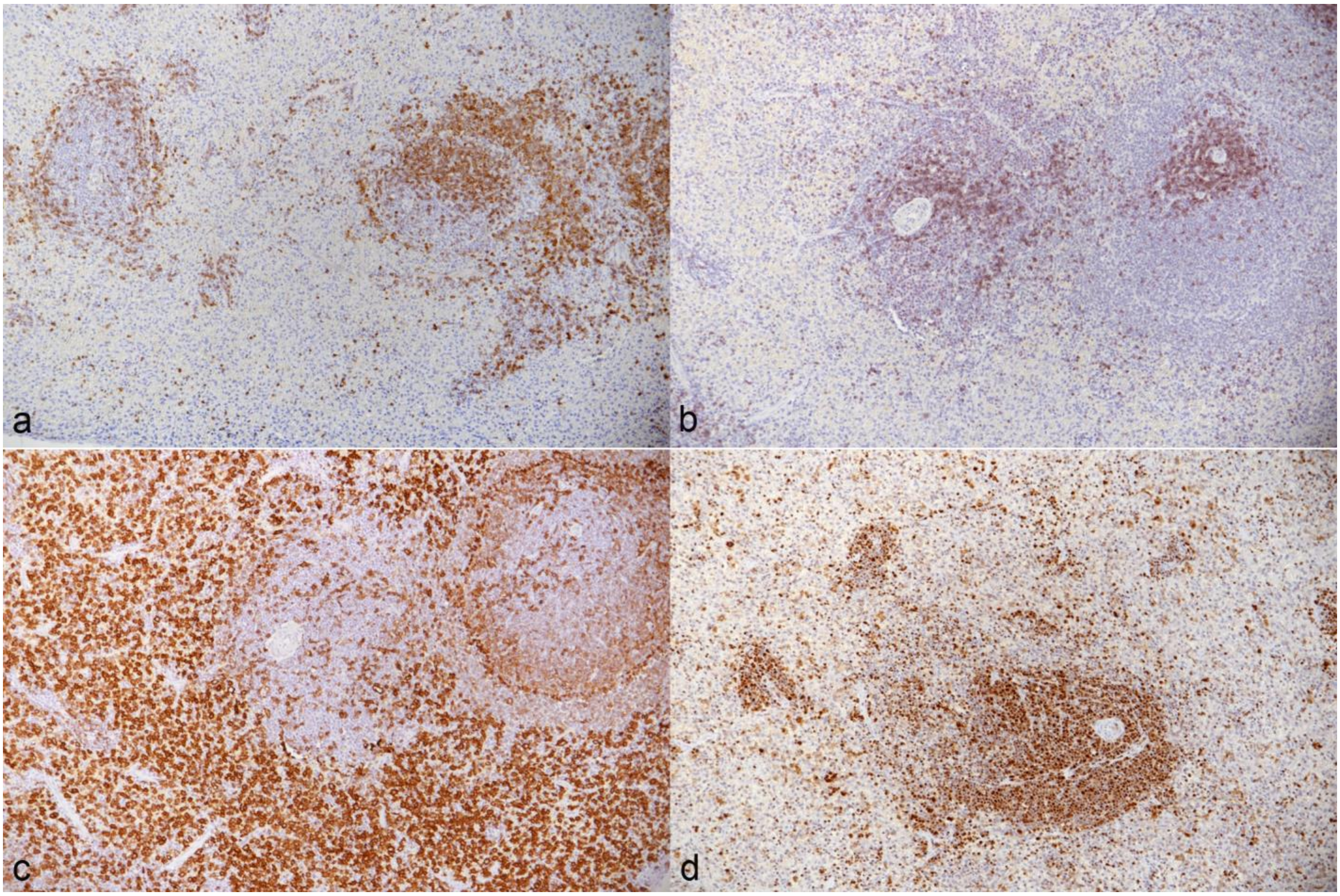


**Figure S6**. Spleen, control animal. **a.** Staining for B cells (CD79a+) identifies small follicles. Immunohistochemistry (IH), hematoxylin counterstain. **b.** Staining for T cells (CD3+) highlights small T cell zones. T cells are present in low numbers in the red pulp. IH, hematoxylin counterstain. **c.** The red pulp contains abundant monocytes/macrophages (Iba1+). IH, hematoxylin counterstain. **d.** Staining for PCNA highlights abundant proliferating cells in the white pulp; they are present in low numbers in the red pulp. IH, hematoxylin counterstain.

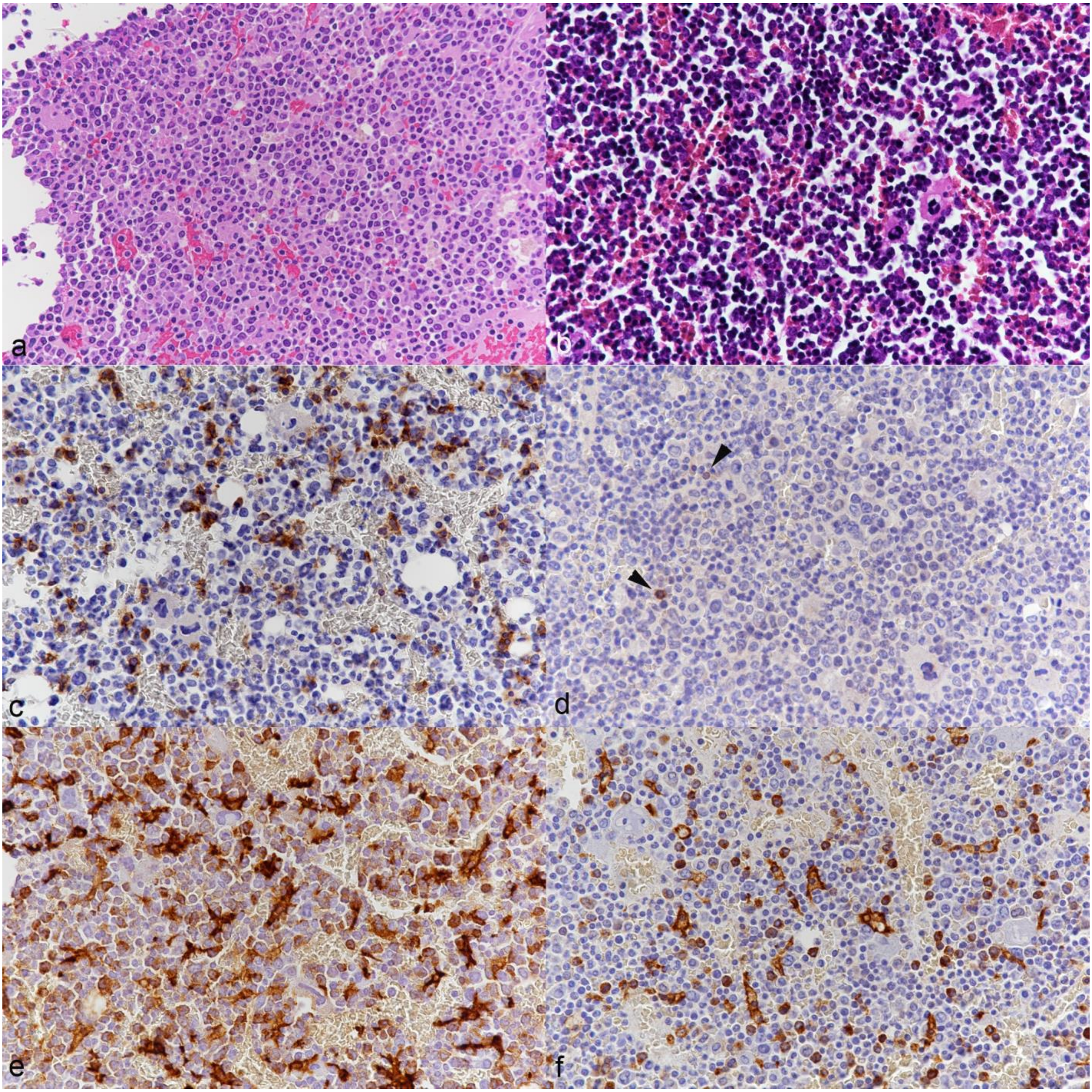


**Figure S7**. Femoral bone marrow. Left column (a, c, e): animal No. 8; right column (b, d, f): control animal. **a, b.** HE stain. The bone marrow is cell rich. Overall, there appear to be more blastoid cells in the infected animals (a) than in the control animal (b). **c, d.** Immunohistochemistry (IH) for T cells (CD3+), hematoxylin counterstain. In the infected animal, there are numerous individual or small groups of T cells (c); in the control animal (d), T cells (arrowheads) are rare. **e, f.** IH for Iba1, hematoxylin counterstain. Numerous positive cells are seen disseminated in the infected animal (e); in the control animals, the number of positive cells is far lower (f).
